# Phosphatidylinositol Cycle Disruption is Central to Atypical Hemolytic-Uremic Syndrome Caused by Diacylglycerol Kinase Epsilon Deficiency

**DOI:** 10.1101/633867

**Authors:** Vincent So, Jing Wu, Alexis Traynor-Kaplan, Christopher Choy, Richard Epand, Roberto Botelho, Mathieu Lemaire

## Abstract

**Background:** Loss-of-function mutations in diacylglycerol kinase epsilon (*DGKE*) cause a rare form of atypical hemolytic-uremic syndrome (aHUS) for which there is no treatment besides kidney transplantation. Highly expressed in kidney endothelial cells, DGKE is a lipid kinase that phosphorylates diacylglycerol (DAG) to phosphatic acid (PA). Specifically, DGKE’s preferred substrate is 38:4-DAG, that is DAG containing stearic acid (18:0) and arachidonic acid (20:4). DAG is produced when phosphatidylinositol 4,5-*bis*phosphate (PtdIns(4,5)P_2_) is cleaved by phospholipase C (PLC). A better understanding of how DGKE deficiency impacts the endothelial lipid landscape is critical to developing a treatment for this condition.

**Methods:** We used orthogonal methods to compare the lipid levels in two novel models of *DGKE* deficiency to their respective controls: an immortalized human umbilical vein endothelial cell (iHUVEC) engineered with CRISPR/Cas9 and a blood outgrowth endothelial cell (BOEC) from an affected patient. Methods included mass spectrometry lipidomics, radiolabeling of phosphoinositides with [^3^H]*myo*-inositol, and live-tracking of a transfected fluorescent PtdIns(4,5)P_2_ biosensor.

**Results:** Unexpectedly, mass spectrometry lipidomics data revealed that high 38:4-DAG was not observed in the two *DGKE*-deficient models. Instead, a reduction in 38:4-PtdIns(4,5)P_2_ was the major abnormality.These results were confirmed with the other two methods in *DGKE*-deficient iHUVEC.

**Conclusion:** Reduced 38:4-PtdIns(4,5)P_2_—but not increased 38:4-DAG—is likely to be key to the pro-thrombotic phenotype exhibited by patients with *DGKE* aHUS.

**TRANSLATIONAL STATEMENT:** Mutations in *DGKE* cause a severe renal thrombotic microangiopathy that affects young children and leads to end-stage renal disease before adulthood. DGKE preferentially phosphorylates diacylglycerol to its corresponding phosphatidic acid (PA), which is then used to synthesize PtdIns(4,5)P_2_ via the phosphatidylinositol cycle. Understanding the disease pathophysiology is necessary to develop a treatment to prevent this outcome. This paper describes how we applied mass spectrometry lipidomics to two novel models of *DGKE* deficiency to investigate how this defect impacts the levels of diacylglycerol, PA and related phosphoinositides in endothelia. Unexpectedly, our data show that the critical abnormality caused by *DGKE* deficiency is not high diacylglycerol, but rather low PtdIns(4,5)P_2_. Restoring endothelial PtdIns(4,5)P_2_ homeostasis may be the cornerstone to treat these patients.

## INTRODUCTION

aHUS is a rare disease that causes renal failure because of thrombosis in kidney glomeruli.^1^ Most genetic forms of aHUS are caused by mutations that lead to hyperactivation of the complement system.^2^ We recently showed that loss-of-function mutations in *DGKE* cause a novel form of aHUS that does not respond to anti-complement therapy.^3^ Typically, an infant with recurrent episodes of renal thrombosis, usually preceded by minor infections, develops end-stage renal disease within 10-20 years of diagnosis.^3, 4^

DGKE is a lipid kinase that phosphorylates DAG to PA; DAG is produced when PLC hydrolyzes PtdIns(4,5)P_2_.^5^ DGKE is unique amongst DGKs because its preferred substrate is 38:4-DAG, a form of DAG containing stearoyl (18:0) and arachidonoyl (20:4) moieties.^6–8^ DGKE is highly expressed in glomerular podocytes and endothelial cells (ECs).^3^ However, of these two cell types, only the latter actively regulates thrombosis.^9^

Based on its enzymatic mode of action, it is generally assumed that *DGKE* deficiency leads to accumulation of its substrate, DAG, and concomitant depletion of the reaction product, PA.^10^ In that context, enhanced DAG-dependent signaling is deemed central to the pathophysiology of *DGKE* aHUS.^10^ While this assumption is eminently reasonable, there are no studies assessing directly how DGKE deficiency impacts the levels of DAG, PA or other related phospholipids in ECs or kidney glomeruli. The high DAG hypothesis thus remains unproven.

To address this issue, we developed a mass spectrometry lipidomics approach to quantify all the relevant lipids from the same sample. We also engineered two novel experimental models of *DGKE* deficiency: *1) DGKE*^**−/−**^ iHUVEC clones modified using Cas9 genome editing, and 2) BOEC derived from a patient with *DGKE* aHUS. Our results, which do not support the high DAG hypothesis but instead point to more complex effects on phosphoinositide biosynthesis, will help guide the development of novel therapies for *DGKE* aHUS, while also providing a unique opportunity to study fatty acyl chain-specific lipid biology.

## CONCISE METHODS

### *DGKE*^−/−^ immortalized HUVEC (iHUVEC)

We used Cas9 RNA-guided nuclease genome editing coupled with dual single-guide RNAs to generate an EC line devoid of DGKE activity. The cell line used is RF24, an immortalized cell line derived from HUVEC (a generous gift from Dr. Ingbord Klaassen, University of Amsterdam).^11, 12^ We used dual Cas9 RNA-guided nuclease genome editing to trigger either a large (14 kb) deletion and/or small indels in the *DGKE* gene (**Supplemental Figure 1**).^13^ This approach yielded three clones with bi-allelic loss-of-function variations by triggering either a macrodeletion within the DGKE gene and/or frameshift mutations. Three wild-type clones were also generated as controls. Cell from passage 4-12 were used for experiments. We used established protocols to measure DGKE mRNA by RT-qPCR^14, 15^ and EC cell surface markers by flow cytometry (CD31, CD54, CD105 and CD144).^16^ All cell lines were systematically tested for the presence of mycoplasma contamination every year with the MycoAlert PLUS Mycoplasma Detection Kit (cat# LT07-705, Lonza, Basel, CHE).

### Human BOEC from a patient with DGKE aHUS

Patient-derived BOEC are complementary to the iHUVEC approach because these cells are not immortalized and they harbor homozygous missense *DGKE* mutations known to cause disease. We used the established protocol to generate BOEC from a patient with molecularly confirmed diagnosis of *DGKE* aHUS (BOEC*DGKE*) and an unrelated adult control subject (**Supplemental Figure 2**).^17^ All participants signed an informed consent form detailing this procedure. BOECs from passages 4-12 were used for experiments. *DGKE* mRNA levels and EC cell surface markers were investigated as described for iHUVEC.

### Mass Spectrometry-Based Lipidomics on EC lines

To normalize the lipidomics data, half of a dish with confluent ECs was processed to quantify protein concentration using the Pierce BCA Protein Assay Kit (Thermo Fisher Scientific) according to the manufacturer’s protocol. The other half was used for acidic lipid extraction with TCA precipitation following an established protocol.^18^ Lipid analytical internal standards for DAG, PtdIns, PtdInsP_2_ and PtdInsP_3_ were added to the TCA precipitates (all from Avanti Polar Lipids, Inc). Extracted lipids were dried under a stream of N_2_ gas and then stored at −80°C. Prior to analysis, phosphate negative charges on phosphoinositides were neutralized by methylation.

Samples were analyzed with mass spectrometry in the multiple reaction monitoring mode, using electrospray and positive ion mode, as described.^18^ Peak areas for acyl-chain specific lipid species and standards were quantified by integrating mass spectrometry curves as previously described.^18^ Peak areas were normalized to total protein levels and to the peak area of the corresponding synthetic lipid standard to account for differential loss of samples during lipid extractions and methylation. For analysis of lipid species without a corresponding synthetic lipid standard, the values were normalized to total protein and expressed as normalized peak areas. It is important to note that our study is optimized to confidently assess the levels of 38:4-containing lipids. While levels for the other measurable lipid species are also provided, the level of confidence should be assumed to be lower because of multiple testing issues.

### Measurement of Phosphoinositides (PtdIns) Using Radioactive ^3^H-inositol Labelling

iHUVEC cells were grown as described above except that just before the cells reached confluence, the growth medium was changed to an inositol-free EC medium with endothelial growth supplements (both products from ScienCell). The solution was supplemented with 75 mCi/mL myo-[2-^3^H(N)] inositol (Perkin Elmer) and 5% dialyzed inositol-free FBS. After 24 hours, adherent cells were scraped and processed following a protocol recently described.^19, 20^ Briefly, lipids were extracted after precipitation and de-acylation. ^3^H-inositol loading was measured with flow scintillation (^3^H window) before storage at −20°C until ready for further processing. Data for specific phosphatidylinositide species were obtained with high performance liquid chromatography (HPLC) followed by flow scintillation based on their known retention times as described.^19^ For each species, we recorded the peak area (total counts), subtracted the background signal, and normalized the resulting data against the parental PtdIns species. Results are expressed as n-fold change in *DGKE*^**−/−**^ compared to control.

### Measuring PtdIns(4,5)P_2_ Using Fluorescently Labelled Lipid Binding Probes

Near-confluent iHUVEC were transfected with plasmids encoding the fluorescent PtdIns(4,5)P_2_ biosensor PH-PLCδ-GFP.^21^ It encodes for the pleckstrin homology (PH) domain of phospholipase C delta (PLCδ) fused to GFP (gift from Dr S. Grinstein).^21^ Cells were analyzed by confocal microscopy 24-48 hours post-transfection and after short exposure to the nuclear stain Hoechst® 33342 (Thermo Fisher Scientific). Data were captured and analyzed using Volocity Software (Perkin Elmer, Woodbridge, Canada). The mean GFP fluorescence intensity per pixel was measured in the plasma membrane (PM) and cytosol (cyt); background signal was subtracted from each value. The semi-quantitative measure of plasmalemmal PtdIns(4,5)P_2_ was quantified as the ratio of PM/Cyt GFP fluorescence.

### Statistical Analysis

All data represent three independent experiments unless otherwise indicated. Data is represented as mean ± standard deviation, unless otherwise stated. All experiments were assessed using unpaired t-tests performed with Prism 7 (GraphPad Software, San Diego, CA).

## RESULTS

### Generation of a *DGKE*-deficient endothelial model

To better understand the pathogenesis of *DGKE* aHUS, we studied the impact of complete *DGKE* deficiency on the lipid profile of ECs. We focused our studies on these cells because they have an established role in regulating thrombosis, whereas podocytes do not.^22^ To this end, we applied CRISPR/ Cas9 to iHUVEC^11, 12^ to generate three *DGKE*^**−/−**^ clones (Figure 1A-B). *DGKE*^**−/−**^ cells have reduced levels of *DGKE* mRNA and protein, are free of mutations at predicted off-target sites, have normal levels of EC surface markers by flow cytometry, and their wound healing ability is similar to that of control cells (**Supplemental Figure 1**).

**Figure 1:**
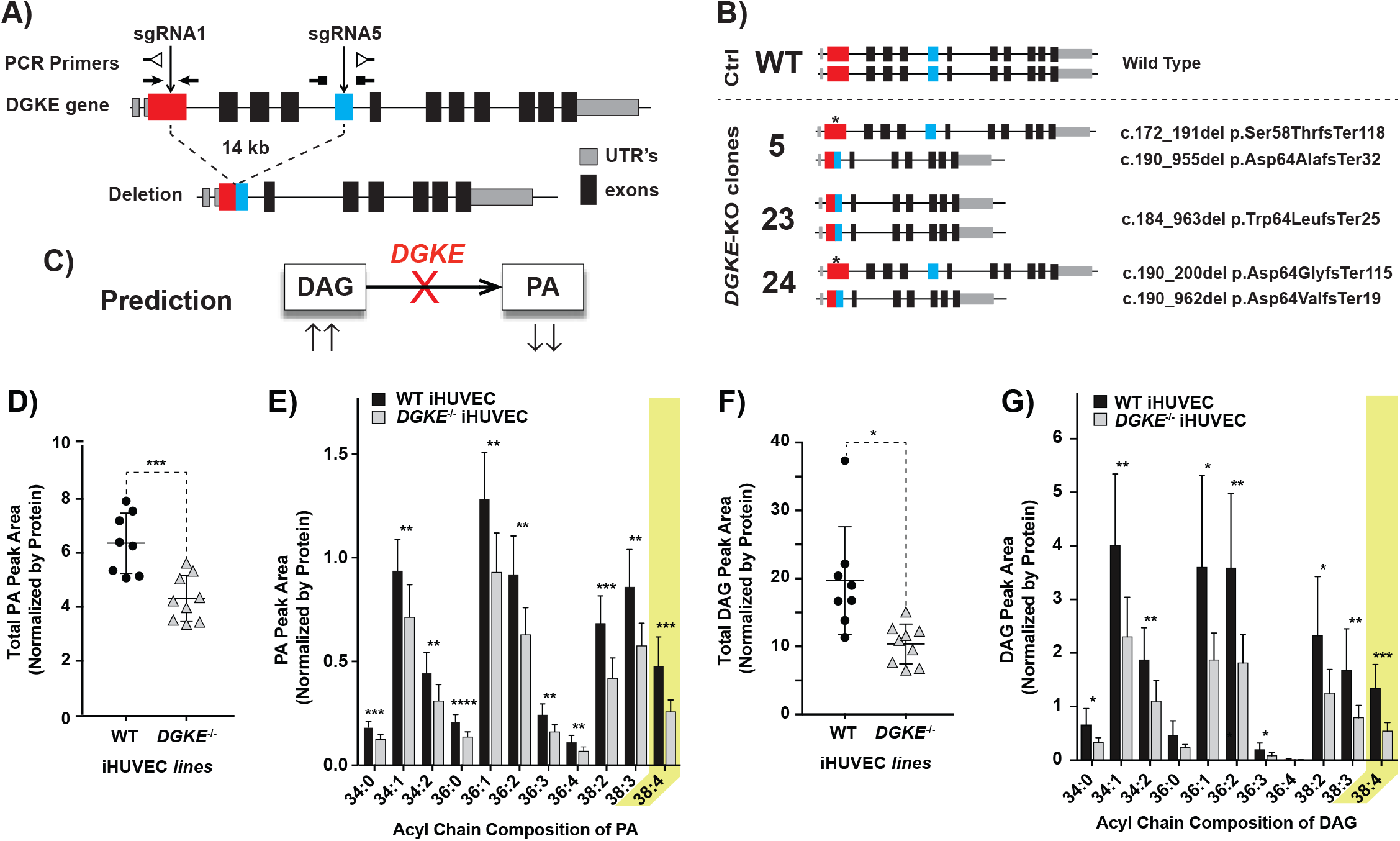
Immortalized *DGKE*^−/−^ iHUVECs have reduced 38:4-PA and 38:4-DAG. A) Strategy to generate *DGKE*^−/−^ endothelial cell lines from immortalized HUVEC (iHUVEC; also known as RF24 cell lines). The relative positions of the sgRNAs used to generate a 14 kilobases (kb) macrodeletion (or small indel) by targeting exons 2 (red) and 6 (blue) within the human *DGKE* gene. The position of the oligonucleotide primers used to amplify the various segments are indicated: closed arrowheads indicate amplifications over short distances (small indels) and open head arrowheads indicate amplification over long segments (macrodeletions). B) The genomic structure and the nucleotide sequences for each *DGKE*^−/−^ clone, derived from PCR and Sanger sequencing. * indicates site where a small indel was created that induces a frameshift. C) Illustration of the enzymatic reaction catalyzed by diacylglycerol kinase epsilon (DGKE). D-G) *DGKE*^−/−^ iHUVEC samples analyzed using mass spectrometry lipidomics to derive the sum total of all measurable PAs (D), the levels for each distinct measurable PA species (E), the sum total of all measurable DAGs (F), the levels for each distinct measurable DAG species (G). Lipids were precipitated from confluent monolayers of *DGKE*^−/−^ or control iHUVEC and used in lipid extractions and methylation, coupled to mass spectrometry-based lipidomics for the endogenous detection of PA and DAG. All experiments were done with cells at steady-state (i.e., not stimulated nor starved). Bars represent the mean levels of each DAG and PA lipid species from at least three independent experiments for each of the three wild-type and 3 *DGKE*^−/−^ clones (6 cell lines in total, and 17 independent measurements). The yellow colored rectangle indicate the values for our main target lipid, 38:4-PA and 38:4-DAG. Error bars represent standard deviation of the mean. * denotes p<0.05, ** p<0.01, *** p<0.001 and **** p<0.0001.

### *DGKE*-deficient iHUVEC have low PA

Since DGKE phosphorylates DAG to PA, the absence of a functional enzyme may be predicted to cause PA reduction (Figure 1C). We compared the lipid composition of wild-type and *DGKE*^**−/−**^ iHUVECs by mass spectrometry to validate this prediction. The sum of all measurable PA species was significantly lower in *DGKE*^**−/−**^ when compared to wild-type (Figure 1D). DGKE is known to preferentially phosphorylate DAG decorated with 38:4 fatty acyl chains –stearic acid, 18:0; arachidonic acid, 20:4– to the corresponding PA. A more detailed lipidomic analysis was therefore undertaken to investigate whether this PA species was particularly affected. As expected, we found that 38:4-PA levels were markedly reduced in *DGKE*^**−/−**^ iHUVEC (Figure 1E). However, PAs decorated with other acyl chain combinations also appeared to be significantly lower in these cells (Figure 1E).

These data confirm that DGKE activity is indeed critical to generate 38:4-PA, a finding consistent with earlier *in vitro* studies on the enzymatic activity of DGKE.^7, 8, 23, 24^ Our results also suggest that the other nine mammalian DGKs –none of which have specificity for 38:4-DAG^25, 26^– cannot fully compensate for *DGKE* deficiency.

### DAG is not elevated in *DGKE*^−/−^ iHUVEC

The reduced formation of PA observed in the *DGKE*-deficient cells should, in principle, be accompanied by accumulation of its precursor, DAG (Figure 1C). Strikingly, we found the total DAG content to be markedly reduced in *DGKE*-deficient iHUVEC (Figure 1F). Detailed mass spectrometric analyses revealed that the decrease was most significant for 38:4-DAG, although other species were also depressed (Figure 1G). These unexpected results suggest that the cellular changes accompanying *DGKE* deficiency are more complex than anticipated.

### Generation of patient-derived EC model of *DGKE* deficiency

The unexpected findings described above may reflect a peculiarity of gene-edited iHUVEC. It was therefore important to validate their occurrence in alternative experimental models of *DGKE* deficiency. Towards this goal, we generated BOEC^17^ from a patient with a novel homozygous *DGKE* mutation (c.A494G; p.D165G; Figures 2A-B) and from an unrelated control. The aforementioned patient displayed classic features of *DGKE* aHUS (**Supplemental Figure 2A**). After confirming the genotypes of the BOEC*DGKE* (Figure 2C), we documented that when compared to controls, these cells have reduced levels of *DGKE* mRNA but similar levels of key EC surface markers (**Supplemental Figures 2B-C**).

**Figure 2:**
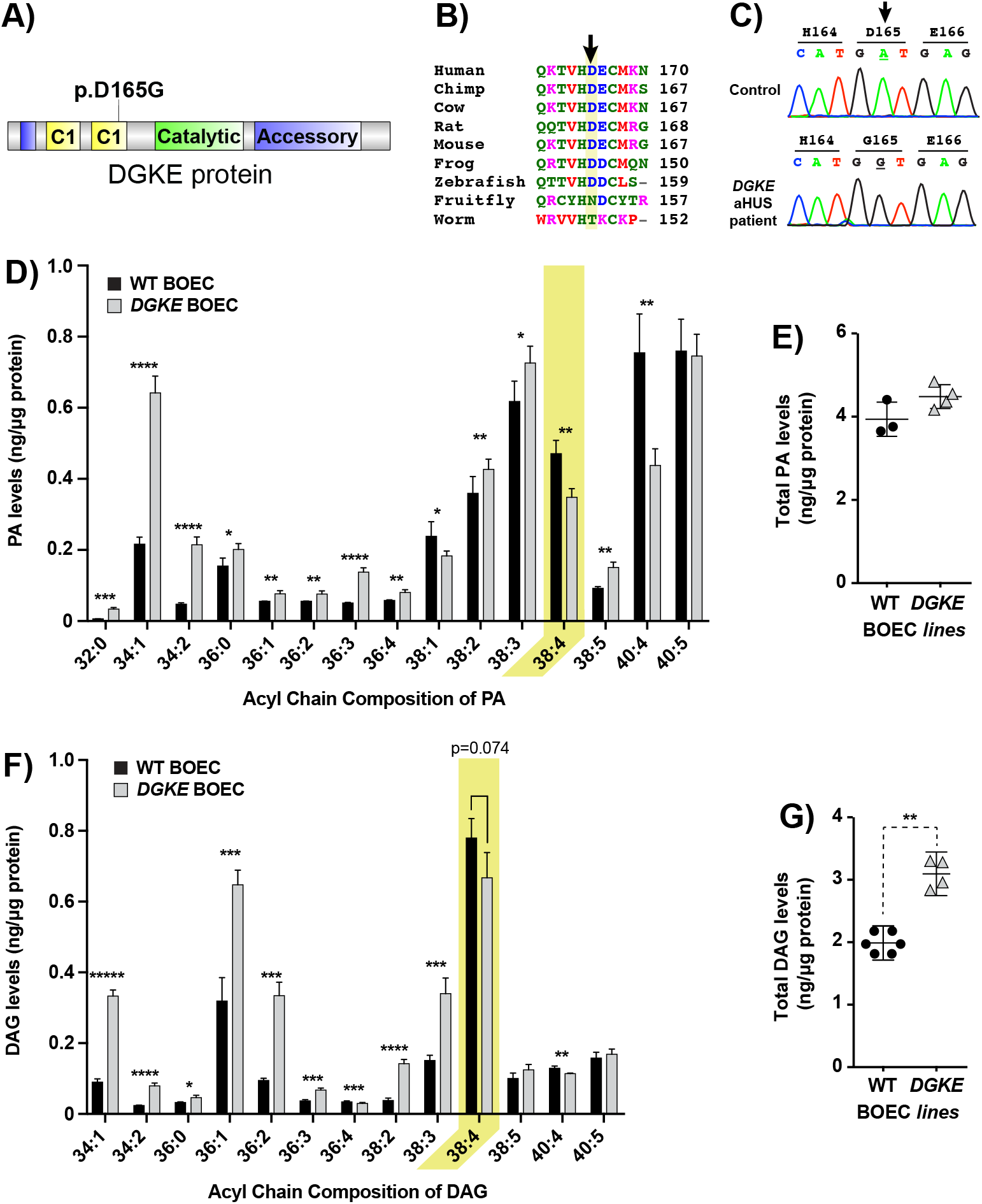
Blood outgrowth endothelial cells (BOEC) from a patient with *DGKE* aHUS have reduced 38:4-PA and 38:4-DAG. A) Illustration of the functional domain of DGKE protein, with the relative position of the missense mutations found in the patient, located in the second C1 domain. B) Alignment of the amino acid sequence of DGKE proteins from a variety of species showing that it is well-conserved down to the zebra fish. C) DNA chromatograms showing the homozygous mutation found in the BOEC derived from the patient sample, c.A494G; p.D165G. Also shown is the sequence amplified from the same genomic segment of control BOEC. D-G) *DGKE* BOEC samples analyzed using mass spectrometry lipidomics to derive the levels for each distinct measurable species of PA (D) or DAG (F) and the sum total of all measurable PAs (E) or DAGs (G). The same protocol described for iHUVEC was used. Bars represent the mean levels of each PA and DAG lipid species from 4 separate experiments with *DGKE* BOEC and 3 with control BOEC. The yellow colored rectangle indicate the values for our main target lipid, 38:4-PA and 38:4-DAG. Error bars represent standard deviation of the mean. * denotes p<0.05, ** p<0.01, *** p<0.001, **** p<0.0001 and ***** p<0.00001.

### 38:4-PA is low and 38:4-DAG is not high in BOEC^*DGKE*^

We used mass spectrometry to assess if PA reductions were also observed in BOEC*DGKE*. Detailed analysis of the discrete PA composition showed that these cells had significantly lower 38:4-PA while having elevated levels for nearly all other PA species measured (Figure 2D). As a result, total PA was marginally higher when comparing mutant to control BOEC (Figure 2E).

We then sought to determine if the level of 38:4-DAG in BOEC*DGKE* would be elevated, conforming to the prediction (Figure 1C), or would instead replicate the unexpected results obtained with *DGKE*^**−/−**^ iHUVEC (Figure 1G). Similar to the pattern observed for PA in BOEC*DGKE*, we found a trend towards lower 38:4-DAG (p=0.074) accompanied by significant increases in most other DAG species (Figure 2F). As a result, total DAG levels were significantly elevated in BOEC*DGKE* (Figure 2G).

Altogether, the parallel reductions in 38:4-PA and 38:4-DAG observed in BOEC*DGKE* resembles that observed with *DGKE*^**−/−**^ iHUVEC, despite the fact that the total PA and DAG data are different between experimental models. The compensatory increases in PA and DAG noted in BOEC*DGKE*, which involved various species decorated with the same acyl chains, suggest the coordinated activation of enzyme sets that are unique to BOEC.

### Low 38:4-PtdInsP_2_ is a core feature of both *DGKE*-deficient experimental models

The reduced 38:4-PA levels observed in both models are attributable to the absence of DGKE activity, but must also be due in part to the reduced availability of its substrate, 38:4-DAG. The paucity of 38:4-DAG, while unexpected, may play an important role in the pathogenesis of *DGKE* aHUS. We therefore proceeded to investigate the mechanism underlying this paradoxical finding. Reduced 38:4-DAG could result from decreased 38:4-PtdInsP_2_, its immediate precursor, to reduced phospholipase C activity, and/or to enhanced activity of other DAG metabolizing enzymes (**Supplemental Figure 3**). That low 38:4-PtdInsP_2_ may be responsible for the abnormal DAG content is suggested by prior studies^8, 27^ that ascribed to DGKE a key role in the phosphatidylinositol (PI) cycle; biosynthesis of PtdIns, the precursor of PtdInsP_2_, requires PA, which is in turn generated from DAG by DGK (Figure 3A).

**Figure 3:**
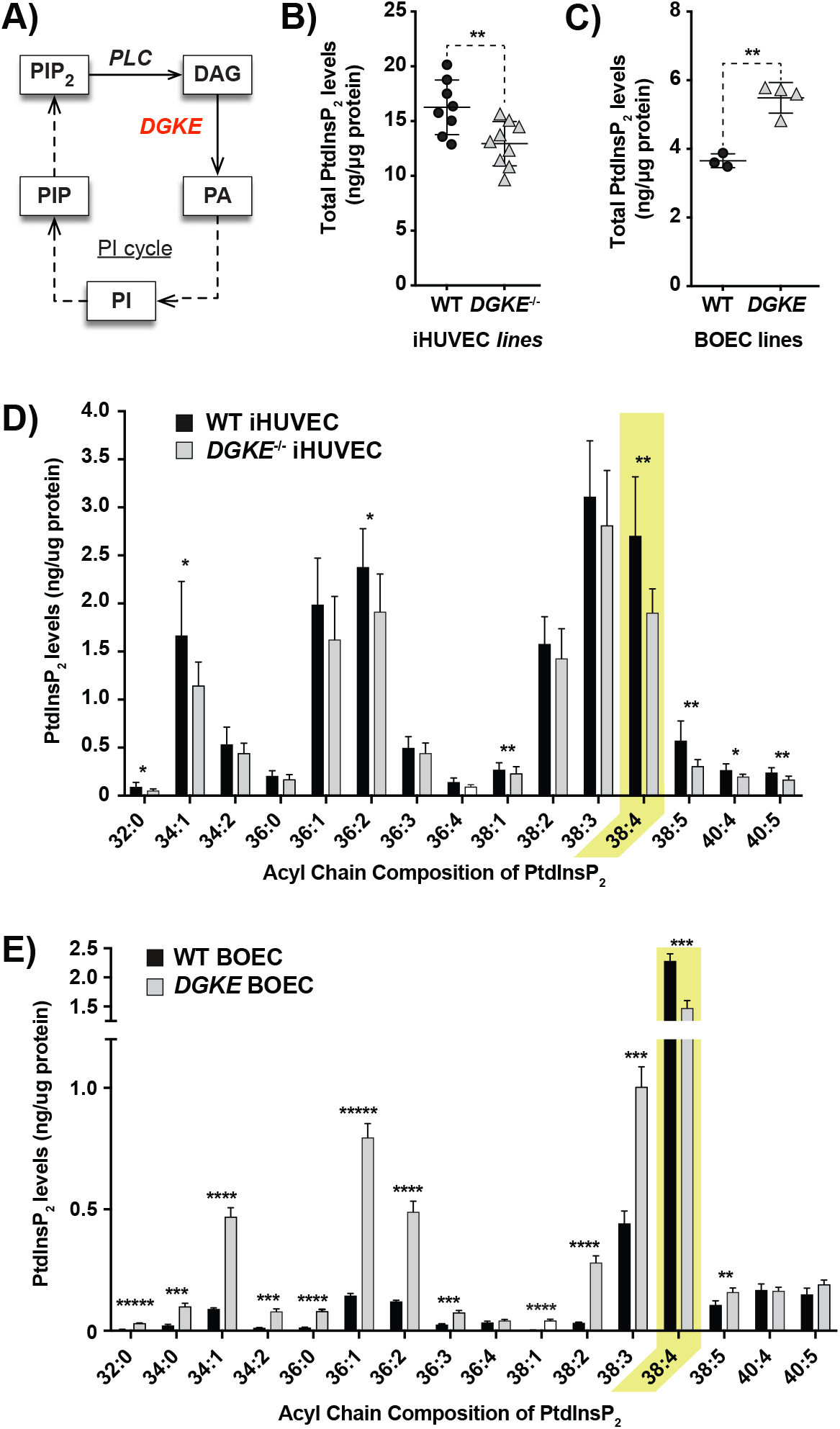
Both experimental models of DGKE deficiency all have reduced 38:4-PIP_2_ levels. A) Illustration of the major enzymes (phospholipase C (PLC) and DGKE) and lipids (PI, PIP, PIP_2_, DAG and PA) implicated in the phosphoinositide (PI) cycle. B-C) The sum total of all measurable PIP_2_ for experiments done with iHUVEC (B) or BOEC (C). Wild-type and *DGKE*-deficient model systems are represented by “Δ” and 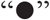, respectively. D-E) The levels for each distinct measurable species of PIP_2_ for experiments done with iHUVEC (D) or BOEC (E). Wild-type and *DGKE*^−/−^ model systems are represented by black and white bars, respectively. The same protocol described for iHUVEC was used except that an internal PIP_2_ standard decorated with 17:0/17:0 was used to improve the reliability of the semi-quantitative measurements. Bars represent the mean levels of each PIP_2_ lipid species from at least three independent experiments (for details on the number of experiments for each model, please refer to Figures 1–3). The yellow colored rectangle indicate the values for the main target lipid, 38:4-PIP_2_. Error bars represent standard deviation of the mean. * denotes p<0.05, ** p<0.01, *** p<0.001, **** p<0.0001 and ***** p<0.00001.

We therefore compared the PtdInsP_2_ content of wild-type versus *DGKE*-deficient cells by mass spectrometry, using the two systems described above. Total PtdInsP_2_ measurements showed significant differences between wild-type and *DGKE*-deficient samples in both systems (Figures 3B-D). As was the case for total DAG, total PtdInsP_2_ content was lower in the *DGKE*^**−/−**^ iHUVEC when compared to wild-type samples. The situation was reversed for BOEC*DGKE*, with slightly higher PtdInsP_2_ content, resembling the findings obtained for DAG (Figure 2G). The similarities between systems were greater when individual PtdInsP_2_ species were analyzed. The content of 38:4-PtdInsP_2_ was reduced in both models of *DGKE* deficiency (Figures 3E-G). Moreover, the net increase in PtdInsP_2_ content recorded in BOEC*DGKE* was attributable to an elevation in PtdInsP_2_ species with the same acyl chain combinations observed for PA and DAG.

Collectively, these results strongly suggest that, by reducing the availability of 38:4-PA, *DGKE* deficiency alters the PI cycle, preferentially affecting the synthesis of 38:4-PtdInsP_2_, the most biologically active form of PtdInsP_2_.^28^

### Reduction in 38:4-PtdInsP_2_ is due to low PtdIns(4,5)P_2_

Mass spectrometric analyses can quantify the content of PtdInsP_2_ with specific acyl chains but cannot distinguish among the three constituent isomers, namely PtdIns(4,5)P_2_, PtdIns(3,4)P_2_ and PtdIns(3,5)P_2_. PtdIns(4,5)P_2_ is generally more abundant than the two other species, and is therefore likely to account for the observed changes, but direct demonstration of this assumption was lacking. Two approaches were used to address this possibility. First we took advantage of the genetically-encoded biosensor PLCδ-PH-GFP that specifically binds to the headgroup of PtdIns(4,5)P_2_.^21^ When expressed in iHUVEC, this probe partitions preferentially to the plasma membrane (Figure 4A), as described for other cell types.^21^ The ratio of plasmalemmal-to-cytoplasmic GFP fluorescence was used as a measure of relative PtdIns(4,5)P_2_ abundance, thereby accounting for differences in transfection efficiency between cells. When estimated in this fashion, the PtdIns(4,5)P_2_ content of all 3 *DGKE*^**−/−**^ iHUVEC clones was 20-30% lower than controls (Figure 4B; Supplemental Figure 4).

**Figure 4:**
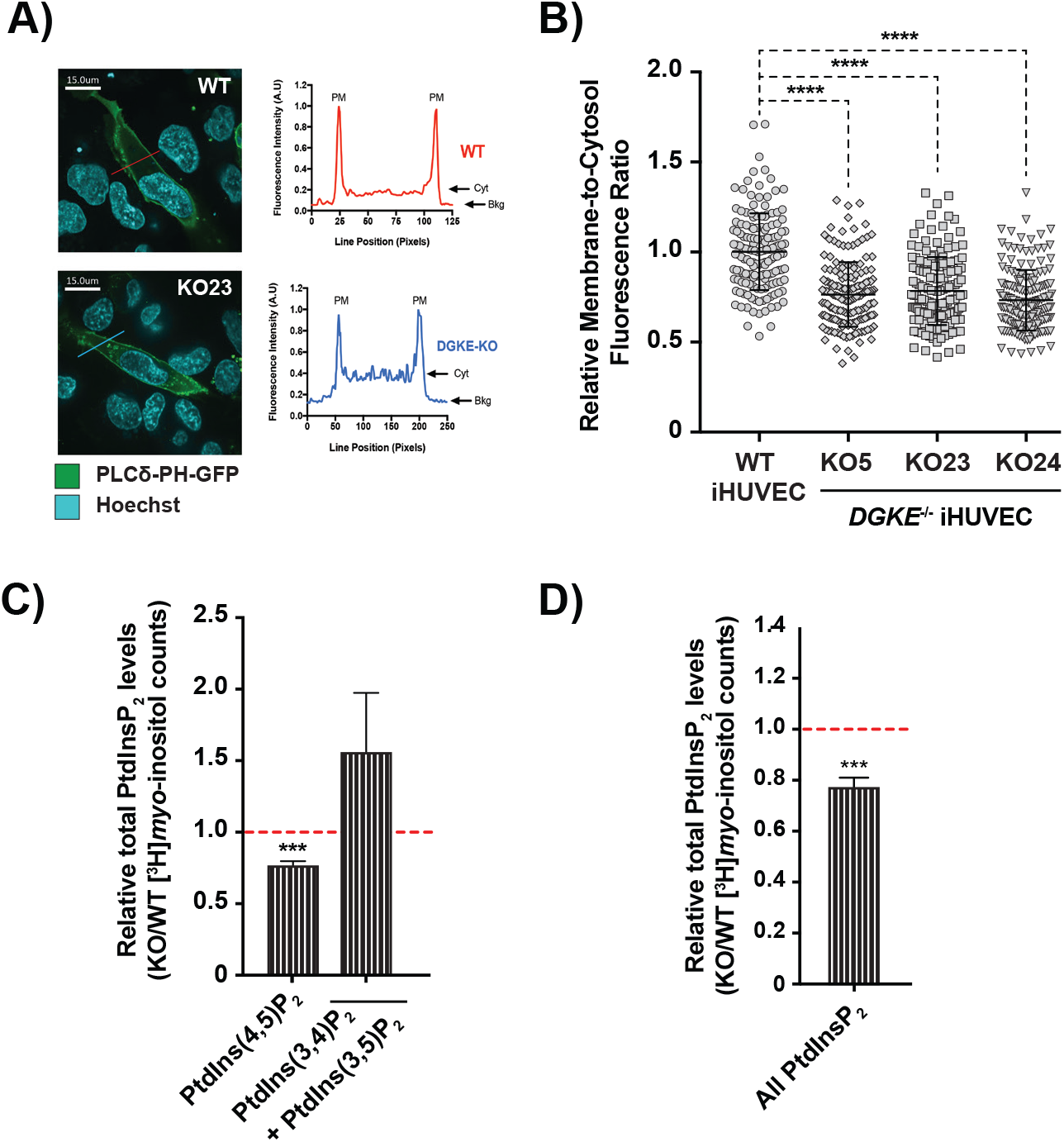
Low 38:4-PtdInsP_2_ is caused by reduced PtdIns(4,5)P_2_ levels in *DGKE*^−/−^ iHUVEC. A) Representative images of cells transfected with GFP-tagged PLCδ-PH lipid binding probe from WT iHUVEC cell line and DGKE-KO iHUVEC (clones KO5, KO23, and KO24). The KO23 clone is displayed as the representative image. Images are representative of 3 independent experiments. Next to each representative image is a line scan plot corresponding to the line drawn across the cell bodies (red for WT, and blue for KO23). The fluorescence intensity in arbitrary units (AU) was normalized to the highest fluorescence value. B) Estimation of the relative levels of plasma membrane PtdIns(4,5)P_2_ after transfection of iHUVEC with a fluorescent PtdIns (4,5)P_2_ biosensor (PLCδ-PH-GFP). The ratio of plasma membrane fluorescence-to-cytoplasm (PM/Cyt) fluorescence was measured for at least 50 cells in three separate experiments for wild-type or *DGKE*^−/−^ iHUVEC. Data represent the mean from aggregated data of at least 150 cells per clone, each point being the value of one cell. The ratio was normalized to the average ratio of PM/Cyt fluorescence across three experiments with wild-type cells. Error bars denote standard deviations. Unpaired t-tests were performed to compare wild-type cells to each knockout clone (KO5, KO23, KO24). C) Relative levels of PtdInsP_2_ isomers using [H]*myo*-inositol metabolic cell labeling followed by liquid scintillation counting of the HPLC eluent. All PtdInsP_**2**_ species measurements were normalized to total PI levels. Values of *DGKE*^−/−^ iHUVEC are expressed as radioactive counts relative to control iHUVEC counts. The dotted red line indicates the expected values if *DGKE*^−/−^ iHUVEC have levels similar to control cells. D) “All PtdInsP_**2**_” data are derived from the sum of radioactive signals of PtdIns(3,4)P_**2**_, PtdIns(3,5)P_**2**_ and PtdIns(4,5)P_**2**_. Data represent the mean of 3 independent experiments. The error bars denote standard deviation of the mean. * denotes p<0.05, *** p<0.001 and **** p<0.0001.

It is challenging to measure PtdIns(3,4)P_2_ or PtdIns(3,5)P_2_ using fluorescent lipid-binding probes because these species are comparatively scarce in cells.^29, 30^ Furthermore, a high-avidity probe for PtdIns(3,4)P_2_ was not available until very recently^31^, and the validity of the only available probe for PtdIns(3,5)P_2_^32^ has been questioned.^33^ To circumvent these problems, we attempted to fractionate the different PtdInsP_2_ isomers by HPLC to resolve the relative abundance. Newly synthesized phosphoinositides were metabolically labeled with [^3^H]*myo*-inositol after incubating confluent cells in inositol-free medium. Lipid extracts were deacylated and head-groups were separated by HPLC and detected by flow scintillation.

Because each experiment requires large amounts of [^3^H]*myo*-inositol, only a single *DGKE*^**−/−**^iHUVEC clone (KO23) was compared with a control. We found that [^3^H]*myo*-inositol uptake was similar between these cell. Reassuringly, newly synthesized PtdIns(4,5)P_2_ levels were significantly decreased (by ≈20%) in *DGKE*^**−/−**^ compared to control iHUVEC (Figure 4B). Of note, the sum of PtdIns(3,4)P_2_ and PtdIns(3,5)P_2_, which could not be independently resolved by the chromatographic scheme applied, was higher in *DGKE*^**−/−**^ cells, although the increase did not attain statistical significance. This increase is unlikely to be of biological significance since the contribution of PtdIns(3,4)P_2_ and PtdIns(3,5)P_2_ to the total PtdInsP_2_ pool is negligible. Indeed, despite the increase in these two species, the total PtdInsP_2_ content –which is dominated by PtdIns(4,5)P_2_– was lower in *DGKE*^**−/−**^ compared to control iHUVEC (Figure 4C).

Overall, the data from 3 orthogonal methods strongly suggest that at least in iHUVEC, *DGKE* deficiency causes a significant reduction in PtdInsP_2_ levels that is mostly explained by abnormal PtdIns(4,5)P_2_ metabolism. This phenomenon is attributable at least in part by changes in 38:4-PtdIns(4,5)P_2_ and is seemingly accompanied by a small compensatory increase in PtdIns(3,4)P_2_ and PtdIns(3,5)P_2_ of unknown physiologic significance.

### Impact of *DGKE* deficiency on PtdIns(3,4,5)P_3_

Finally, we sought to determine if the reduced 38:4-PtdInsP_2_ found in *DGKE*^**−/−**^ iHUVEC translated into lower 38:4-PtdIns(3,4,5)P_3_ levels. This would be an important finding because *DGKE* deficiency would then simultaneously affect two major signaling pathways: one dependent on PLC, and the other on phosphatidylinositol 3-kinase (PI3K; Figure 5A). We obtained robust and reproducible mass spectrometry lipidomics data even though PtdIns(3,4,5)P_3_ is considerably less abundant than PtdInsP_2_ in cells in basal state (not serum starved). We found that only 38:4-PtdIns(3,4,5)P_3_ levels were significantly lower in *DGKE*^**−/−**^ iHUVEC, compared to control cells (Figure 5B). None of the other species was significantly affected. Using the same approach, we also found trends towards lower 38:4-PtdIns(3,4,5)P_3_ levels in BOEC*DGKE*, but this difference was not statistically significant (Figure 5C).

**Figure 5:**
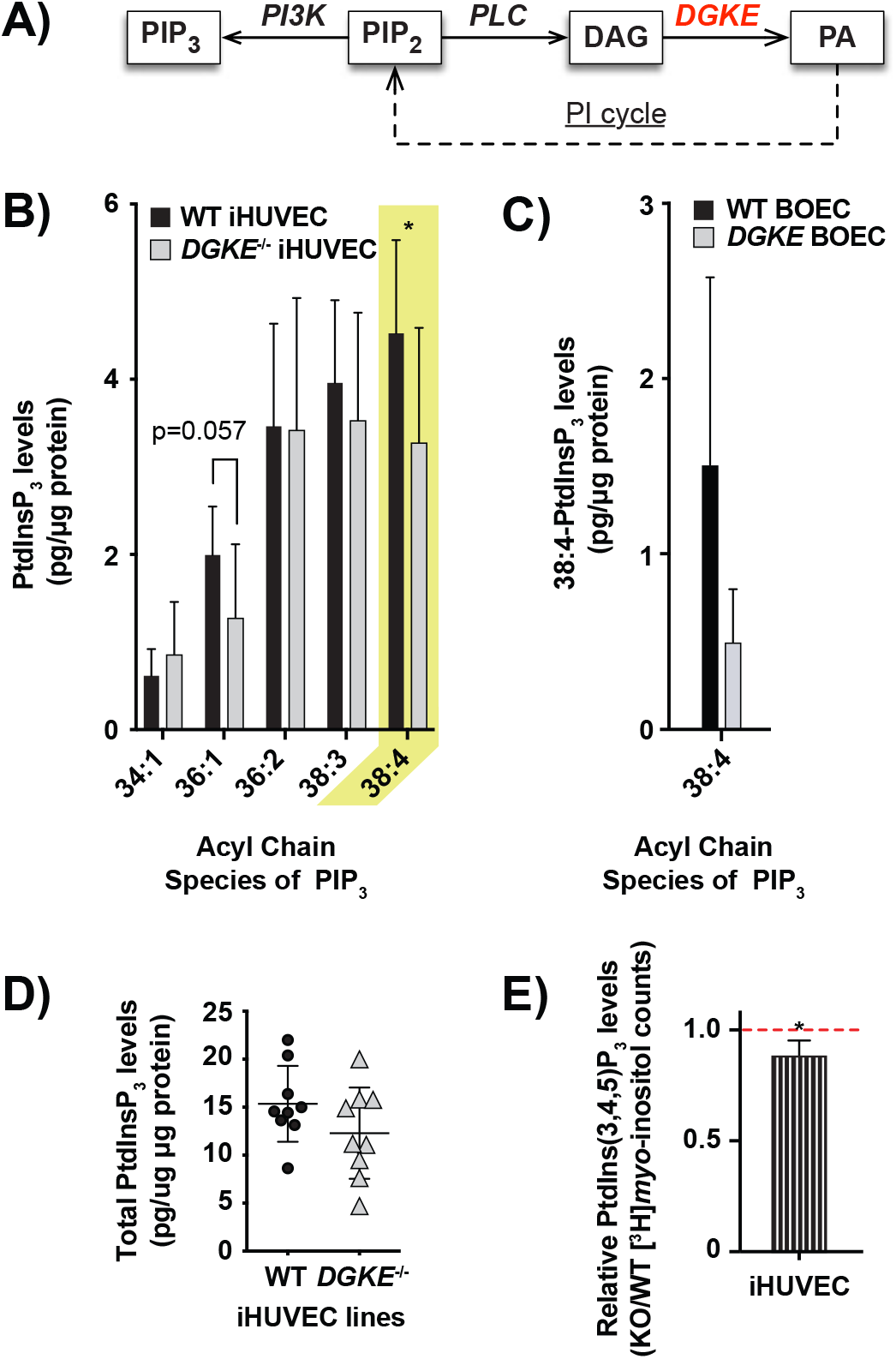
PtdInsP_3_ data for both experimental models of *DGKE* deficiency: measurements by lipidomics and radioactive inositol labeling. A) The pathways responsible for metabolizing PtdIns(4,5)P_2_ to the two main bioactive metabolites, DAG and PtdIns(3,4,5)P_3_. The PI cycle, which is responsible for synthesizing PtdIns(4,5)P_2_ from PA, is also illustrated. B-C) The levels for each distinct measurable species of PtdInsP_3_ for experiments done with iHUVEC (B) or BOEC (C), analyzed using mass spectrometry lipidomics. The yellow colored rectangle indicate the values for the main target lipid, 38:4-PtdInsP_3_. An internal PtdInsP_3_ standard decorated with 17:0/17:0 was used to improve the reliability of the semi-quantitative measurements. Bars represent the mean levels of each PtdInsP_3_ lipid species from at least three independent experiments (for details on the number of experiments for each model, please refer to Figures 1–3). D) The sum total of all measurable PtdInsP_3_ for mass spectrometry lipidomics experiments done with iHUVEC (p=0.157). E) Relative levels of PtdIns(3,4,5)P_3_ using [^3^H]*myo*-inositol metabolic cell labeling followed by liquid scintillation counting of the HPLC eluent as described for PtdInsP_2_ isomers. PtdIns(3,4,5)P_3_ measurements were normalized to total PI levels. Values of *DGKE*^−/−^ iHUVEC are expressed as radioactive counts relative to control iHUVEC counts. The dotted red line indicates the expected values if *DGKE*^−/−^ iHUVEC have levels similar to control cells. Error bars represent standard deviation of the mean. * denotes p<0.05.

In *DGKE*^**−/−**^ iHUVEC, the sum of all PtdIns(3,4,5)P_3_ isomers was only marginally lower in *DGKE*-deficient cells (Figure 5D). However, the difference was significant when newly synthesized PtdIns(3,4,5)P_3_ was assessed by HPLC analysis of cells radiolabelled with [^3^H]*myo*-inositol (Figure 5E).This suggests that the pool of 38:4-PtdIns(3,4,5)P_3_, and its precursor 38:4-PtdIns(4,5)P_2_, turns over faster than other species. Taken together, these data suggest that reduced 38:4-PtdIns(4,5)P_2_ content associated with *DGKE* deficiency causes a secondary reduction in 38:4-PtdIns(3,4,5)P_3_ levels.

## DISCUSSION

We often presume the root cause of *DGKE* aHUS to be excessive DAG-dependent signaling (e.g., via protein kinase C (PKC)).^10, 34^ However, it is unclear if it elevates DAG in affected tissues of patients. Our findings are inconsistent with this notion. We showed that while 38:4-PA was low in *DGKE*-deficient cells, the content of 38:4-DAG was not increased concomitantly. We obtained similar results in two distinct experimental models of *DGKE* deficiency. Instead of the predicted DAG elevation, the salient and most consistent finding was the paucity of 38:4-PtdIns(4,5)P_2_ (Figures 3 & 4). We, therefore, conclude that DGKE is essential for the generation 38:4-PA, which is itself required for the PI cycle to generate and maintain plasmalemmal 38:4-PtdIns(4,5)P_2_ levels. We suggest that *DGKE* deficiency should be re-defined as an inborn error of phosphoinositide biogenesis.

The low levels of 38:4-PtdIns(4,5)P_2_ are likely a direct consequence of reduced 38:4-PA, the critical substrate to initiate the PI cycle. Since 38:4-PtdIns(4,5)P_2_ accounts for ~50-80% of PtdIns(4,5)P species in primary cells and fresh tissue^18, 35–37^, it is likely central to many cellular processes, including signal transduction, membrane transport and actin cytoskeleton remodeling.^5^ *In vivo*, these functions are carried out directly by 38:4-PtdIns(4,5)P_2_, or indirectly via its hydrolysis to 38:4-DAG and Ins(1,4,5)P_3_ by PLC, or by phosphorylation to 38:4-PtdIns(3,4,5)P_3_ by PI3K.^5^ While PLC and PI3K isoforms have no selectivity for PtdIns(4,5)P_2_ species, 38:4-PtdIns(4,5)P_2_ is their main substrate given its preponderance in cells. The remarkable phenotype associated with *DGKE* deficiency suggests that kidneys are uniquely sensitive to reduced levels of 38:4-PtdIns(4,5)P_2_ or its products.

It is striking that low 38:4-PtdIns(4,5)P_2_ is also the primary abnormality of cells deficient in other enzymes with 38:4 specificity, namely lysophosphatidylinositol-acyltransferase (LPIAT1)^38^, lysocardiolipin acyltransferase (LYCAT)^19, 39^ and CDP-DAG synthase 2 (CDS2).^40^ Interestingly, CDS2 silencing in Zebrafish and HUVEC primarily disrupts endothelial functions.^41^ While cells invest vast resources to produce 38:4-PtdIns(4,5)P_2_, little is known about the functional importance of various acyl chains. Studying *DGKE* deficiency provides a unique opportunity to investigate this question.

As stated above, *DGKE*-deficient ECs did not have high 38:4-DAG levels when compared to controls. This unexpected observation may be due to reduced PLCδ or PLCγ activation, two major PLC isoforms that are expressed in endothelial cells^42, 43^ and use PtdIns(4,5)P_2_ as their preferred substrate. The PH domain of PLCγ that is instrumental in its recruitment to the membrane, selectively binds PtdIns(3,4,5)P_3_^44^, which is depleted *pari passu* with PtdIns(4,5)P_2_ in *DGKE*-deficient cells. The activation of PLCδ is similarly regulated, but via the binding of its PH domain to PtdIns(4,5)P_2_ itself.^45,46^

Given our results, it is prudent to re-interpret prior studies that assumed that elevated DAG is central to *DGKE* deficiency. The increased p38 and p44/42 activation observed earlier^34^ following *DGKE* siRNA knockdown in HUVEC may not be attributable to high PKC activity. Accordingly, PKC activity was not increased in these cells.^34^ In this context, it will be important to assess if *DGKE* silencing phenocopies *DGKE* knockout in terms of 38:4-PtdIns(4,5)P_2_ levels, in which case the depletion of this phosphoinositide (or of a metabolite produced therefrom) may explain the altered activity of the two kinases. This is important because the phenotype of siRNA knockdown is not always concordant with that of a gene knockout.^47–49^ Lastly, it is noteworthy that our results are in agreement with earlier studies that measured lipid levels in brain extracts^50^ and fibroblast cell lines^23^ derived from the original *Dgke*^**−/−**^ and control mice. While we noted no differences in DAG levels, both reports highlighted the reduced PtdInsP_2_.^23, 50^

That overexpressing DGKE in porcine aortic ECs triggered a reduction in 38:4-DAG^51^ may at first appear to contradict our results. However, a likely explanation relates to the essentiality, in both scenarios, of the physiological rate-limiting step of the PI cycle which, judging from our results, is the generation of 38:4-PtdIns(4,5)P_2_. Excessive heterologous expression of DGKE should reduce the levels of its substrate DAG regardless of 38:4-PtdIns(4,5)P_2_ levels. In contrast, *DGKE*-deficient cells are exquisitely sensitive to the rate-limiting step since 38:4-PtdIns(4,5)P_2_ is in short supply.

Our findings show that therapies aimed at reducing DAG levels are unlikely to benefit patients with *DGKE* aHUS. We should instead focus efforts on restoring 38:4-PtdIns(4,5)P_2_ levels, perhaps with intravenous infusion of the lipid itself or, more likely, of its more membrane-permeant precursors. This could mitigate the predicted effects on the actin cytoskeleton, the suboptimal anti-thrombotic protein levels (via defective exocytosis) or decreased nitric oxide production (via poor response to shear stress). Repurposing glycerol^52, 53^ may be an attractive candidate therapy since its uptake and phosphorylation –two key steps in *de novo* PA synthesis^54^– are markedly increased in *Dgke*^**−/−**^ cells.^55^ The propensity of patients to develop renal thrombosis following minor infections^4^ suggests that glomerular lipids help prevent excessive thromboinflammation.^56^ Acute dampening of systemic inflammation with immunomodulatory agents could thus provide alternative treatment options.

## Supporting information

Detailed methods

## ACKNOWLEDGMENTS

We thank Dr. Sergio Grinstein (Cell Biology Program, SickKids Research Institute) for helpful discussion and comments on the manuscript. VS was supported by the Frederick Banting and Charles Best Canada Graduate Scholarships (CGS-M) and the RESTRACOMP Master’s Scholarship from the Hospital for Sick Children. RJB is funded by Natural Sciences and Engineering Research Council, Canada Research Chair Program, Early Researcher Award and Ryerson University. ML is supported by New Investigator Awards from KRESCENT/CHIR and CCHCSP. This research is funded by an operating grant from KRESCENT/CIHR.

