## Supplemental Figures for "Phosphatidylinositol Cycle Disruption is Central to Atypical Hemolytic-Uremic Syndrome Caused by Diacylglycerol Kinase Epsilon Deficiency"

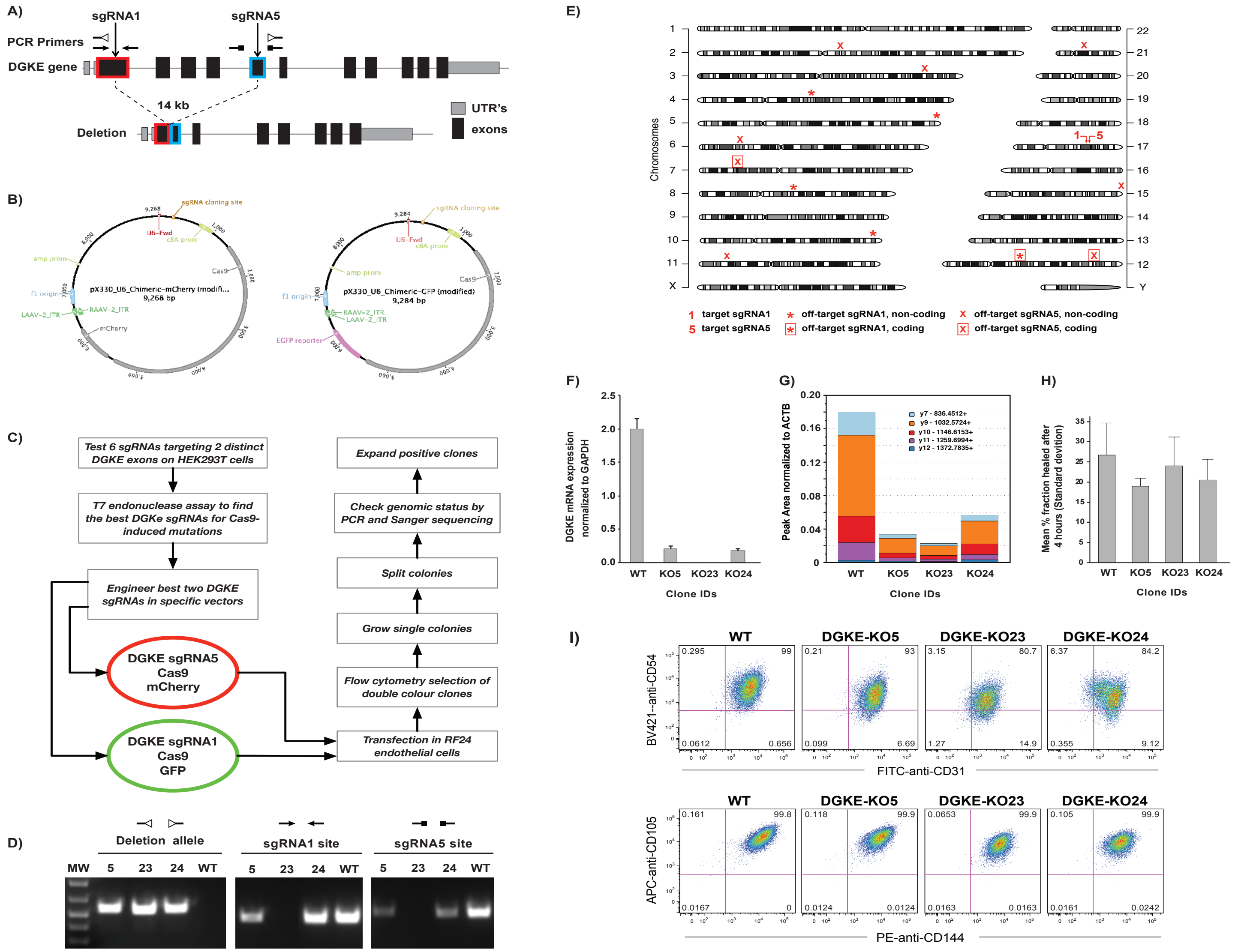


### Supplementary figure 1: Generation of *DGKE*$^{-/-}$ immortalized HUVECs (RF24).

A) Illustration of the relative position of the binding sites for the two single guide RNA (sgRNAs) designed to target two distant coding exons of the *DGKE* gene to generate a 14 kilobases deletion, or coding frameshift mutations in either exons. The relative positions of the PCR primers used in panel C are indicated.

B) The structure of the two plasmids used to transfect Cas9 and the sgRNAs and either of two fluorescent markers into the cells of interest.

C) The workflow used to generate the *DGKE*$^{-/-}$ iHUVEC clones.

D) PCR products observed when probing with primers designed to reveal the presence of the macrodeletion and sequencing for indels.

E) The three *DGKE*$^{-/-}$ iHUVEC clones were screened for all loci predicted to be coding or non-coding off-targets sites for either of the two sgRNAs. The positions of these loci are indicted on the chromosome illustrations, along with the two target sites (labeled 1 and 5).

F) RT-qPCR was used to demonstrate that the three *DGKE*$^{-/-}$ iHUVEC clones have negligible DGKE mRNA expression when compared to wild-type cells, after normalizing to GAPDH levels. Similar results were obtained when DGKE mRNA levels were normalized to ACTB levels (data not shown).

G) Targeted mass spectrometry proteomics was used to show that the DGKE proteins are significantly lower in the three *DGKE*$^{-/-}$ iHUVEC clones when compared to wild-type cells (levels shown relative to those of ACTB).

H) Scratch assay was used to show that the three *DGKE*$^{-/-}$ iHUVEC clones have wound healing capacity that is not significantly different from that of wild type cells.

I) Flow cytometry analysis of the three *DGKE*$^{-/-}$ iHUVEC clones show that they express the four endothelial-specific cell surface markers (CD31, CD54, CD105 and CD144) at levels similar to that of wild type cells.

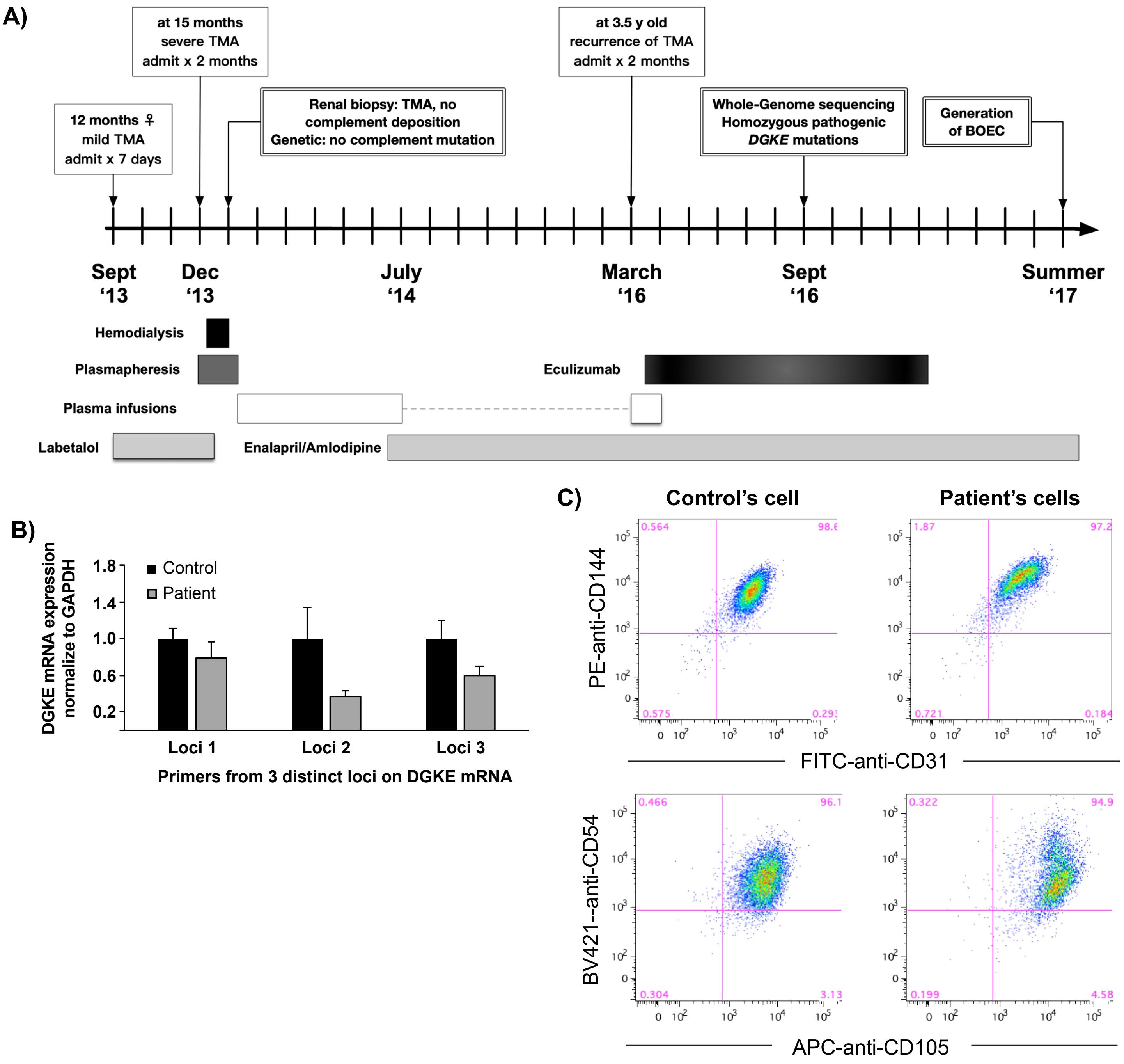


**Supplementary figure 2: Generation of BOEC from a patient with confirmed diagnosis of *DGKE aHUS*.**
A) Clinical course of the patient with *DGKE* p.D165G showing multiple disease recurrences before age 5. Key diagnostic events are shown in box with double stroke contour. Various treatments instituted are illustrated at the bottom of the timeline. The patients remains stable on amlodipine/enalapril with fluctuating levels of proteinuria.
B) RT-qPCR was done to assess the level of DGKE mRNA expressed in BOEC derived from the patient when compared to controls. mRNA levels from the patient's cells are expressed relative to wild-type BOEC, after normalizing for GAPDH levels. Similar results were obtained when normalized to ACTB levels (data not shown)
C) Flow cytometry analysis of the control and patient-derived BOEC showing that they express the four endothelial-specific cell surface markers (CD31, CD54, CD105 and CD144) at similar levels.

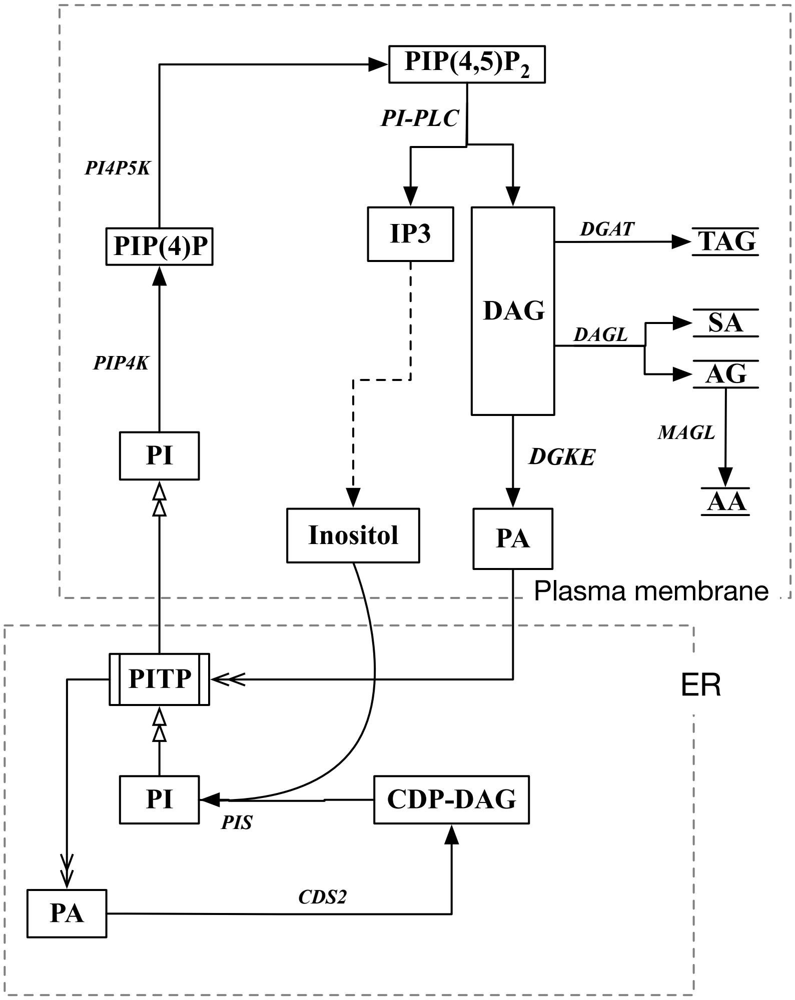


**Supplementary figure 3: DAG metabolism via DGKE, DAG lipase (DAGL) or DAG acyltransferase (DGAT).**
Illustration of the major pathways for the metabolism of DAG produced by phospholipase C (PLC). Also indicated are various members of the PI cycle described in the text.

Other abbreviations used include:
AA, arachidonic acid
AG, arachidonoyl glycerol
CDS2, CDP-DAG synthase 2
IP3, inositol 3-phosphate
MAGL, monoacylglycerol lipase
PI, phosphatidylinositol
PIP, PI 4-phosphate
$PIP_3$, PI 3,4,5-trisphosphate
PI3K, PI 3-kinase
PIP4K, PIP 4-kinase
PI4P5K, PI4P 5-kinase
PIS, PI synthase
PITP, PI transport protein
SA, stearic acid
TAG, triacylglycerol

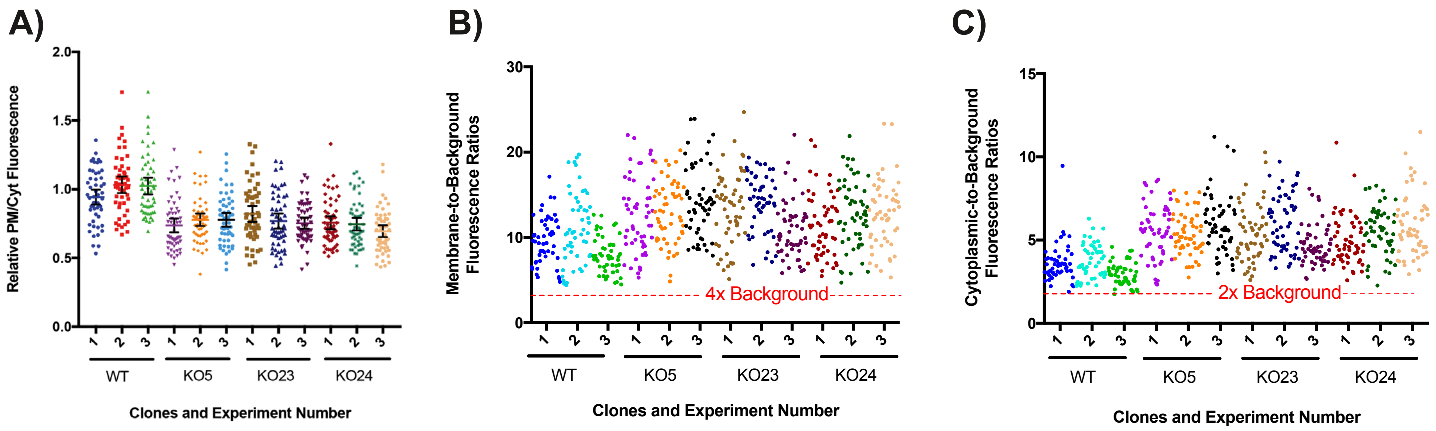


**Supplementary figure 4: Detailed $PIP_2$ Biosensor Fluorescence Measurements.**
A) The ratio of plasma membrane fluorescence-to-cytoplasm ((PM/Cyt) fluorescence for each individual experiment and clone (WT, KO5, KO23, and KO24). Data represents the mean ratio for at least 50 cells (n=50) and each point denotes a value from a single cell. The ratio was normalized to the average ratio of PM/Cyt fluorescence across three WT experiments. Error bars denote standard deviations.
B-C) Illustration of the membrane-to-background and cytoplasm-to-background fluoresence ratios for each data point. Each point represents the ratio of PM/Bkg or Cyt/Bkg for one individual cell. For each individual experiment, at least 50 cells (n=50) were analyzed for each experimental group. iHUVEC cells transfected with the PLCδ-PH-GFP probe were quantified and utilized in subsequent analysis if the membrane fluorescence reached a threshold of 4 times or higher than background, and if the cytoplasmic fluorescence reached a threshold of 2 times or higher of background.

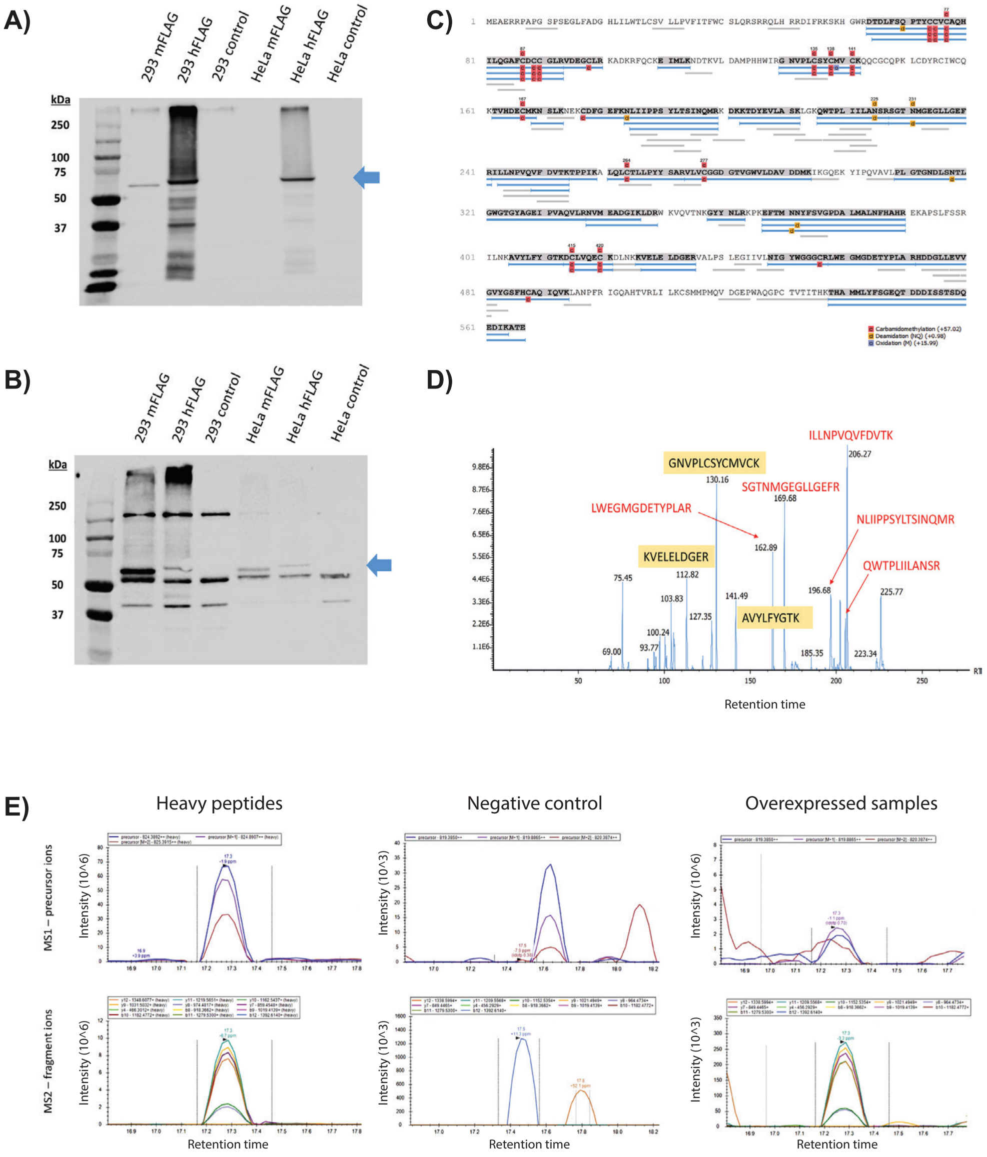


**Supplementary figure 5: Targeted Mass Spectrometry Approach to Detect DGKE Proteins.**
A) Cell lysates overexpressing human DGKE and mouse DGKE in HeLa and HEK293 cell lines were used in western blots to assess DGKE expression using anti-FLAG antibodies. The blue arrows denote the corresponding band for DGKE (~64 kDa).
B) Western blot from overexpressed samples using an anti-DGKE antibody. The blue arrows denote the corresponding band for DGKE (~64 kDa).
C) Sequence alignment of detected DGKE peptide fragments (in blue) from tandem MS/MS from overexpressed samples and DGKE protein sequence.
D) Annotated chromatogram of DGKE protein from overexpressed samples denoting peptide fragments (blue fragments in Panel C) that were highly abundant and matched DGKE protein sequences. Yellow fragments are unique to human DGKE protein, and red fragments are common to mouse and human DGKE protein.
E) Representative images of peak distribution of precursor ions (top panels) and isotopic distribution from fragment ions (bottom panels) for the detection of peptide fragment LWEGMGDETYPLAR. Heavy peptides were utilized as a positive control (left panel). Fragmented DGKE ions from overexpressed DGKE samples had a similar isotopic distribution compared to the heavy labeled peptides, although the intensity of the peaks is much lower (right panel).
