## Supplementary material for "Phosphatidylinositol Cycle Disruption is Central to Atypical Hemolytic-Uremic Syndrome Caused by Diacylglycerol Kinase Epsilon Deficiency": Detailed methods

### TABLE OF CONTENTS

1. Generation, maintenance and characterization of cell lines 2
*1a. Generation of Immortalized DGKE-KO RF24 Cell Line 2*
*1b. Generation of Human Blood Outgrowth Endothelial Cells (BOECs) 3*
*1c. Culture Conditions for Immortalized RF24 Cells and BOECs 4*
*1d. Flow Cytometry for Endothelial Cell Surface Markers 4*
*1e. mRNA isolation and quantification by RT-qPCR 4*
2. Targeted Mass Spectrometry-Based Proteomics for DGKE Protein Quantification 5
*2a. Identification of DGKE Peptides for Targeted Mass Spectrometry 5*
*2b. Overexpression of DGKE in Cell Lines 6*
*2c. Cell Lysis and Sample Preparation 7*
*2d. Protein Digestion and Sample Preparation 7*
*2e. Parallel Reaction Monitoring (PRM) for DGKE Protein Detection 7*
3. Measuring PI Cycle Phospholipids Using Mass Spectrometry-Based Lipidomics 8
*3a. Phase-Contrast Imaging of Cell Confluence 8*
*3b. Cell Lysis and Lipid Precipitation 8*
*3c. Lipid Extraction and Sample Preparation 9*
*3d. Mass Spectrometry Procedures 10*
4. Measurement of Phosphoinositides Using Radioactive $^3$H-inositol Labelling 10
*4a. Culture Conditions and $^3$H-inositol Labelling 10*
*4b. Deacylation and Lipid Extraction 11*
*4c. High Performance Liquid Chromatography (HPLC) 12*
*4d. Radioactive Normalization and Analysis 12*
5. Measuring PI(4,5)$P_2$ Using Fluorescently Labelled Lipid Binding Probes 13
*5a. Culture Conditions and Plasmid Transfection 13*
*5b. Confocal Microscopy 13*
6. Statistical Analysis 13
7. References 13

Tables
- Method Table 1: List of gRNA and potential off-target sites 2
- Method Table 2: Primers to investigate Dgke mRNA expression from human cell lines. 5

### 1. Generation, maintenance and characterization of cell lines

#### *1a. Generation of Immortalized DGKE-KO RF24 Cell Line*

The overall goal was to generate a human endothelial cell line harboring a large multi-exonic deletion in the gene encoding for DGKE. A total of 6 single guide RNAs (sgRNA) targeting two distinct exons of DGKE gene were tested on HEK293 cell lines. These were designed using on line tool from Dr. Zhang lab (http://tools.genome-engineering.org). The most optimal DGKE sgRNAs for CRISPR/Cas9-mediated gene editing was selected using the T7 endonuclease assay. This assay relies on the ability of this enzyme to cleave genomic DNA that contains heteroduplexes within the targeted segment.[1] One sgRNA targeting exon 2 of DGKE gene was inserted in a plasmid that also encoded GFP, while the second sgRNA targeting exon 6 was inserted in a plasmid containing mCherry.[2] Plasmids pX330-U6-Chimeric-GFP and pX330-U6-Chimeric-mCherry were gifts from Dr. Lin Ye (Univeristy of California, San Francisco).[2] The constructs were co-transfected into immortalized HUVECs (iHUVEC; also known as the RF24 cell line)[3,4], aiming to generate a large 14 kilobase (Kb) macrodeletion or short microdeletions at either exon sites. Then, flow cytometry was performed to select for cells expressing both GFP and mCherry. Individual cells were seeded and grown into clonal populations. As controls, three wild-type clones were also generated using the same approach (without gene editing). To check for genome editing, PCR and Sanger sequencing were performed using 3 sets of PCR primers, which would amplify exon 1, exon 5, and across the deletion triggered by cutting at exons 1 and 5. The three clones that were found to be *DGKE*$^{-/-}$ were expanded to be used in subsequent experiments (these cells had either biallelic macrodeletions or a 14 Kb heterozygous deletion accompanied by a small deletion in exon 1 or 5). Off-target sites were checked by Sanger sequecing, with particular emphasis on the loci that could affect coding exons (Method Table 1).

**Method Table 1**: List of gRNA and potential off-target sites

| Single-guide RNAs | Genes targeted (mRNA) | Nucleotide sequences | Loci in human genome (GRCh38/hg38) | Target category (coding) |
|---|---|---|---|---|
| sgRNA1 | *DGKE* (NM_003647) | CAAGCACGGGTGGCGCGACA**CGG** | chr17:56834969 | on-target |
| | *LRP1* (NM_002332) | CAAGCAGGGATGGGGCGAGA**GGG** | chr12:57212901 | off-target |

| | | | | |
|---|---|---|---|---|
| sgRNA5 | *DGKE* (NM_003647) | CAACGATCTATCCAATACAT**TGG** | chr17:56,848,746 | on-target |
| | TMEM116 NM_138341 | CAACATTCTTTCCAATCCAT**TAG** | chr12:111931556 | off-target |
| | CREB5 NM_001011666 | CAACCATATATCCAAGTCAT**AAG** | chr7:28821421 | off-target |
| Notes: There was a total 41 off-target sites for gRNA1, 2 are in genes and 1 is exonic. There was a total of 59 off-target sites for gRNA5, 3 are in genes, and 2 are exonic. Only of exonic off-target sites are listed here. PAM sequences are in bold. | | | | |

#### *1b. Generation of Human Blood Outgrowth Endothelial Cells (BOECs)*

To generate the BOECs, we followed the protocol published by Martin-Ramirez and colleagues in 2012.[5] First, a total of 48 mL of blood was collected from patients or donors into six Sodium Heparin CPT Vacutainer tubes (BD Biosciences, cat# 362753). The tubes were gently inverted 8-10 times to mix the anticoagulant additives with the blood. The tubes were then left at room temperature for 30-60 minutes before remixing and centrifugation at room temperature for 30 minutes at 1,600 × g in a horizontal rotor centrifuge. The serum and mononuclear cell layers were removed and added to 8 mL of 1X Phosphate-buffered saline (PBS) (Wisent Inc., cat# 311-425- CL) supplemented with 10% fetal bovine serum (FBS) in a 50 mL conical tube. The sample was centrifuged at 520 × g for 10 minutes at room temperature. The resulting supernatant was carefully removed before being discarded. The cell-rich pellet was gently resuspended in 10 mL of 1X PBS supplemented with 10% FBS.

Viable cells were then counted using a hemocytometer after staining of a small sample with Trypan Blue Solution (Thermo Fisher Scientific, cat# 15250061), according to the manufacturer's protocol. A cell suspension containing approximately 107 cells/mL cells was prepared using the growth medium specially designed for BOEC (see Section c for details). A volume of 4 mL of the cell suspensions was added to 6-well culture dishes. Each well was pre-coated with x mL of 50 µg/mL collagen IV to improve cell adherence (Thermo Fisher Scientific, cat# 17104019). The cells were incubated in a humidified incubator at 37°C with 5% $CO_2$. After 24 hours, the media was gently replaced with fresh BOEC media, taking precautions to minimize disturbing the cells. The media was replaced daily for 7 days post-seeding, and then replaced every other day until BOECs obtain characteristic cobblestone morphology was observed. When following this protocol, none of the other blood-borne cells that were present originally survive.

#### *1c. Culture Conditions for Immortalized RF24 Cells and BOECs*

Immortalized HUVECs (RF24 Line) and BOECs were maintained in Endothelial Basal Medium (EBM-2; Lonza, cat# CC-3156) prepared in single bottle batches with SingleQuots Kits that contain a propritetary mixture of growth factors, cytokines, and supplements (Lonza, cat# CC-4176) and 5% FBS (Thermo Fisher Scientific, cat# 12483020). Cells were dissociated from culture dishes using TrypLE Express (Thermo Fisher Scientific, cat# 12604-013), recombinant cell-dissociation enzymes with trypsin-like activity. The cells were maintained in a humidified incubator at 37°C with 5% $CO_2$. BOECs were grown on culture dishes pre-coated with 50 µg/mL collagen IV (Thermo Fisher Scientific, cat# 17104019). Immortalized RF24 lines were used from passages 4-20 and BOECs were used from passages 4-12.

#### *1d. Flow Cytometry for Endothelial Cell Surface Markers*

DGKE wild type and null cell lines were analyzed for cell surface marker expression by flow cytometry, using a protocol adapted from that described by Reiss et al., 2004.[6] Briefly, cells grown to confluence in 6-well dishes and retrieved by treatment with TrypLE in PBS (Invitrogen/ Thermo Fisher Scientific, Waltham, MA). The cells were then washed with FACS buffer (containing 1X HBSS; 2% FBS; 10mM HEPES, pH 7.2; 10 mM $NaN_3$) before sequential incubations with the primary antibodies followed by washes with FACS buffer. All antibodies were directly conjugated with fluorochromes. The antibodies used included anti-human CD31- FITC (Cat#MA1-19587, Pierce/Thermo Fisher Scientific, Waltham, MA); anti-human CD54- BV421 (Cat#564077, BD Biosciences, San Diego, CA); anti-human CD105-APC (Cat#17-1057- 41, eBioscience/Thermo Fisher Scientific, Waltham, MA); anti-human CD144-PE (Cat#580410, BD Biosciences, San Diego, CA). 7-AAD was used to eliminate dead cells. The cells were not fixed with paraformaldehyde. Samples were analyzed using the flow cytometer BD LSR II (BD Biosciences, San Diego, CA). Data analysis was performed using the FlowJo software (Mac version 10.4.2).

#### *1e. mRNA isolation and quantification by RT-qPCR*

mRNA from the cell lines was isolated using TRIzol reagent (Cat#15596026, Invitrogen) following manufacturer's instruction. DNase I digestion was performed to avoid genomic DNA contamination in the RNA samples (cat#M0303L, NEB). RNA was further purified using PureLink RNA Mini Kit

(Cat#12183018A, Invitrogen). Reverse transcription (RT) was performed using SuperScript III Reverse Transcriptase (Cat#18080093, Invitrogen) and random hexamers were used as primers. Quantitative PCR (qPCR) was done using Power SYBR Green PCR Master Mix (Cat#4367659, Applied Biosystems) on a StepOne Real-Time PCR system (Cat#4376357, Applied Biosystems). cDNA (20ng) was used as a template for each qPCR reaction. The $2^{-\Delta\Delta Ct}$ method was used to calculate the relative mRNA abundance, after normalization using GAPDH.[7,8] The primers used for these assays as listed in Method Table 2.

**Method Table 2**: Primers to investigate Dgke mRNA expression from human cell lines.

| Target regions | Forward/Reverse Primer sequences | Product size (bp) | Notes on target sites |
|---|---|---|---|
| 1 | F: CAGCGTCGTTCTCCTCCTG<br>R: GCCTCTCCGCTTCCATCTTC | 105 | NM_003647; 5'UTR<br>NM_003647; exon 1 |
| 2 | F: TGGGAGAAGGACTGTTGGGA<br>R: GGGAGAAGAGTACAGAGTTGTAG | 109 | NM_003647; exon 3<br>NM_003647; exon 4 |
| 3 | F: GTCTGCTGGAAGTCGTTGGA<br>R: GCATTGGCATCATGGAGCAC | 132 | NM_003647; exon 9<br>NM_003647; exon 10 |

### 2. Targeted Mass Spectrometry-Based Proteomics for DGKE Protein Quantification

#### *2a. Identification of DGKE Peptides for Targeted Mass Spectrometry*

In a preliminary experiment, we used shotgun Liquid Chromatography Tandem Mass Spectrometry (LC-MS/MS) approach to detect the top 3000-5000 most abundant proteins within the cell, but we were unable to detect DGKE protein in WT RF24 cells or any of the DGKE-KO RF24 cell lines. As a result, we developed a mass spectrometry-based targeted proteomics approach with parallel-reaction monitoring (PRM), as previously described in the literature[12–14], to selectively detect DGKE peptide fragments.

To develop a targeted PRM approach, we first overexpressed FLAG-tagged mouse DGKE (mDGKE) and FLAG-tagged human DGKE (hDGKE) protein in HEK293 and HeLa cell lines (**Supplemental Figure 8A-B**). Cells were transfected using the protocol described in Section 2b. Protein quantifications were done using the protocol described in Section 2c. The overexpressed DGKE samples were in a standard LC-MS/MS experiment to identify proteotypic peptides that are above the threshold for detection by mass spectrometry in our system. Several peptides were identified that aligned with DGKE protein sequences (**Supplemental Figure 8C-D**). Of these peptides, some were specific to mouse DGKE protein, specific to human DGKE protein, or were common between both species. The peptides fragments that were common between both species were synthesized commercially with a heavy Lysine (Lysine-$^{13}C_6$, $^{15}N_2$) or a heavy Arginine (Arginine-$^{13}C_6$, $^{15}N_4$) at the C-terminal of every peptide.

The synthetic heavy peptides were validated by running targeted PRM mass spectrometry for endogenous DGKE peptide fragments. In these experiments, synthetic heavy peptide for DGKE were added to cell lysates overexpressing DGKE immediately prior to mass spectrometry and were used to select and confirm the monitoring of the correct light peptides. Using the fragmented ions of the heavy peptide as a reference, endogenous DGKE peptide fragment ions that displayed corresponding retention time (± 0.1min) and fragmentation (**Supplemental Figure 8E**).

#### *2b. Overexpression of DGKE in Cell Lines*

Stock plasmids containing the human DGKE gene inserted into a p3xFLAG-CMV™-7.1 vector and the empty 3xFLAG-CMV™-7.1 vector was generously provided by the Epand Lab (McMaster University, Hamilton, Ontario). The plasmid was transformed into chemically competent Escherichia coli (E. coli) using heat shock transformation. Briefly, 50 µL of chemically competent E. coli was incubated with 100 ng of stock plasmid DNA on ice for 30 minutes. The mixture was placed in a 37°C water bath for 1 minute, followed by incubation on ice for 2 minutes. Then, mixtures were added to 500 µL of Luria-Bertani (LB) broth and grown at 37°C for 1 hour. Afterwards, 100-200 µL of the reaction was plated onto an LB agar plate supplemented with 100 µg/mL of ampicillin antibiotics to select for ampicillin-resistant colonies. These were inoculated into LB cultures and grown overnight at 37°C on an orbital shaker set at 250 rpm.

E. coli cultures were pelleted using centrifugation at 7,000 × g at 4°C for 10 minutes. The supernatant was discarded, and the pellet was flash frozen on dry ice and stored at -80°C, prior to plasmid isolation. The plasmid was isolated using a HiPure Plasmid Midiprep Kit (Invitrogen, Ref No. K210004) according to manufacturer's protocols and the concentration was determined using a Nanodrop 2000c Spectrophotometer (Thermo Scientific, cat# ND2000) at wavelength 260 and 280 nm to measure DNA concentrations.

Prior to transfection, cells were grown to 60-80% confluence on glass coverslips coated with 50 µg/mL collagen IV (Thermo Fisher Scientific, cat# 17104019) in 12-well culture plates. Transfections were performed using the Lipofectamine LTX with Plus Reagent Kit (Thermo Scientific, cat# 15338100). Briefly, 1 µg of plasmid DNA was incubated with 200 µL Opti-MEM (Thermo Fisher

Scientific, cat# 31985062) and 2.5 µL of Plus Reagent (Thermo Fisher Scientific, cat# 11514015) for 10-15 minutes at room temperature. Following incubation, 3 µL of Lipofectamine reagent was added to the mixture and incubated at room temperature for 30 minutes to allow for formation of the transfection complexes. These were then added to the growth medium overlaying the target cultured cells for 4-6 hours. Growth medium containing untrasfected complexes was aspirated and replaced with fresh growth medium. The cells were grown to confluence and analyzed 24-48 hours post-transfection. Immediately prior to confocal microscopy, cells were incubated with 1µg/mL of Hoechst® 33342 (Thermo Fisher Scientific, cat# 62249) to identify the nuclear region.

***2c. Cell Lysis and Sample Preparation***

Confluent cells were washed once with ice cold 1X PBS. The supernatant was then discarded and the cells were scraped into RIPA lysis buffer containing 25 mM Tris (pH 8.0), 150 mM sodium chloride (NaCl), 1% NP-40, 0.1% sodium dodecyl sulfate (SDS), 1% sodium deoxycholate, and protease inhibitors (Roche, catalog no. 11836153001). Cells were immediately scraped using a plastic scraper. The mixture was incubated on ice for 30-60 minutes with gentle agitation to help promote cell lysis. Protein concentration was quantified using the Pierce BCA Protein Assay Kit (Thermo Fisher Scientific, cat# 23225) according to the manufacturer's protocol.

***2d. Protein Digestion and Sample Preparation***

Cells were lysed as previously described (see Section c). Approximately 1-10 mg of cell lysate was dissolved in 8M urea, 50 mM Tris-HCl (pH= 8.0), 4 mM dithiothreitol (DTT) in a reaction volume up to 1 mL. The sample was then resuspended in 50-100 µL of 50 mM $NH_4HCO_3$ (pH=8.3) and DDT was added up to a final concentration of 10 mM. The sample was heated at 60°C for 30 minutes, and then cooled to room temperature. Fresh iodoacetamide was added to the solution up to a final concentration of 10 mM and incubated at room temperature in the dark for 15 minutes. The iodacetamide was inactivated by adding DTT to a final concentration of 40 mM. The sample was diluted with 50 mM $NH_4HCO_3$ until the urea concentration was below 1M, and the pH was readjusted to 7.5-8.5 if necessary. Modified trypsin (mass spectrometry grade) was added to a final protease: protein ratio of 1:50 to 1:100 (w/w). The samples were digested overnight at 37°C, followed by lyophilization.

***2e. Parallel Reaction Monitoring (PRM) for DGKE Protein Detection***

Digested and lyophilized samples were mixed with 50 fmol of heavy labelled peptides in 0.1%

formic acid buffer solution. Samples were analyzed on a Q-Exactive HF-X (ThermoFisher, San Jose, CA) outfitted with a nanospray source and EASY-nLC nano-LC system (ThermoFisher, San Jose, CA.). Peptide Mixtures were loaded onto a 75µm x 50 cm PepMax RSLC EASY-Spray column filled with 2µM C18 beads (ThermoFisher San, Jose CA) at a pressure of 800 Bar and a temperature of 60C. Peptides were eluted over 30 min at a rate of 250 nl/min using a gradient set up as 0-42% linear gradient (0.1% Formic acid; and Buffer B, 0.1% Formic Acid in 80% acetonitrile). Peptides were introduced by nano-electrospray into the Q-Exactive HF-X mass spectrometer (Thermo-Fisher). The instrument was configured using the preset PRM method option in positive polarity, with the default charge state set to 2. Resolution was 30,000, with an AGC target of 5e5 and a maximum ion time of 22ms. The quadrupole isolation window was set to 0.5 m/z, and a normalized collision energy was set to 27 for each peptide.

**3. Measuring PI Cycle Phospholipids Using Mass Spectrometry-Based Lipidomics**

The protocols described below are modified from previously published work done by Dr. Alexis-Traynor Kaplan f(ATK Innovation, Analytics and Discovery and the University of Washington).[15]

***3a. Phase-Contrast Imaging of Cell Confluence***

Confluent live cells were imaged using an epifluorescence microscope (DMIRE2, Leica Microsystems) equipped with a Hamamatsu Camera Controller (ORCA-ER), and MAC 6000 Shutter Controller (LudI Electronic Products Ltd) due to the fact that confluence has been demonstrated to influence fatty-acyl chain profiles of cellular phosphoinositides.[15]

***3b. Cell Lysis and Lipid Precipitation***

Prior to lipid precipitation, the adherent cells were washed with 1X cold PBS. Half of the confluent dish of cells was scraped using a rubber scraper. Cells in suspension were transferred to a sterile Eppendorf tube to be used for protein normalization. The cell suspension was pelleted by centrifugation at 1,500 × g at 4°C for 10 minutes. The supernatant was discarded and the pellet was re-suspended in 200-300 µL of RIPA lysis buffer containing 25 mM Tris (pH 8.0), 150 mM sodium chloride (NaCl), 1% NP-40, 0.1% sodium dodecyl sulfate (SDS), 1% sodium deoxycholate, and protease inhibitors (Roche, cat# 11836153001). The mixture was incubated on ice for 30-45 minutes with gentle agitation to lyse cells. Protein concentration was quantified using the Pierce BCA Protein Assay Kit (Thermo Fisher Scientific, catalog# 23225) according to the manufacturer's protocol.

In parallel, 1.5 mL of 0.5M ice-cold trichloroacetic acid (TCA) solution was added directly to the cells covering the other half of the dish. After 5 minutes (on ice), an additional 250 μL of 0.5M ice-cold TCA solution was added to the culture dish (total volume 1.75 mL). The cells were then scraped using a plastic cell scraper into a sterile 2 mL Eppendorf tube. The mixture was vortexed at top speed for 30 seconds, and incubated on ice for an additional 5 minutes (step 1). The mixture was then centrifuged at 20,000 x g at 4°C for 5 minutes to pellet the TCA precipitate (step 2). The supernatant was carefully removed and discarded with a micropipette, without disturbing the pellet (step 3). Afterwards, 1 mL of solution containing 5% (w/v) TCA and 10 mM EDTA was added to the pellet (step 4). Steps 1 to 4 described above were repeated in the exact same order twice. Following the second wash with 5% (w/v) TCA and 10 mM EDTA, the supernatant was completely removed, and the final TCA precipitate pellet was stored at -80°C prior to lipid extraction.

***3c. Lipid Extraction and Sample Preparation***

Prior to lipid extraction, lipid analytical internal standards were added to the TCA precipitates. This procedure was used to improve the accuracy of our measurements by injecteing samples with known amounts of lipid analytic standards with an acyl chain composition that do not exist in cells (e.g., 17:0-20:4, or 37:4). The lipid analytical internal standards used were from Avanti Polar Lipids, Inc. The lipid standards used were all from Avanti Polar Lipids Inc except otherwise indicated: 17:0-17:0 DAG [diheptadecanoin] (Nu-Chek Prep, Inc., cat# D-156), 17:0-20:4 PA (cat# LM-1402), 17:0-20:4 PI(4,5)$P_2$ standard (cat# LM-1904), 17:0-20:4 PI(3,4,5)$P_3$ (cat# LM-1906), 17:0-20:4 PI (cat# LM-1502).

An acidic lipid extraction was then performed by adding 670 μL of chloroform:methanol:12N HCl (40:80:1) mixture to the TCA precipitate. The sample was vortexed vigorously for 2-5 minutes and incubated on ice for 10 minutes. Then 650 μL of ice-cold chloroform was added and the sample was vortexed for 2-5 minutesand allowed to sit for 5 min at 4°C. Afterwards, 300 μL of ice cold 1M HCl was added. The sample was then vortexed for 2 minutes and centrifuged at 10,000 × g for 2 min at room temperature. The lower phase was then collected in a new 2 mL microcentrifuge tube. A theoretical lower phase was generated by creating a chloroform:methanol:1.74 M HCl mixture (86:14:1 v/v/v). The theoretical lower phase was added to the upper phase and the mixture was vortexed for 2 minutes, and centrifuged. The lower phase was then combined with the previously collected lower phase. The lower phase was then dried under a stream of $N_2$ gas and was immediately methylated or stored at -80°C.

Following lipid extraction, phosphate negative charges on phosphatidylinositides were neutralized by methylation. For methyl derivatization of phosphate hydroxyls, 90 μL of methanol/$CH_2Cl_2$ mixture (4:5 v/v) was added to the dried lipid extracts. After vigorous vortexing, 20 μL of 2 M trimethylsilyldiazomethane (TMS-DM) was added and the mixture incubated at room temperature for 1 h in sealed microfuge tubes. The samples were once again dried under $N_2$ gas.

#### *3d. Mass Spectrometry Procedures*

The dried samples were resuspended in 20-100 μL of 100% liquid chromatography mass spectrometry grade methanol. An aliquot of the methyl derivatized sample was separated on a C4 column in an acetonitrile formic acid gradient monitored by a Waters XEVO TQ-S MS/MS in multiple reaction monitoring mode (MRM) using electrospray and positive ion mode. The gradient was initiated with 10 mM formic acid in water / 10 mM formic acid in acetonitrile (33:67 v/v), held for 1 min, then increased to 15:85, v/v in 9 min following injection, held at 85% for 2 min and then re-equilibrated to starting conditions for 3 min. Sodium formate (50 μM in water/acetonitrile, 1/1, v/v) was infused into the post-column eluate using the Intellistart Fluidics of the Waters XEVO TQ-S MS/MS to promote formation of positively charged sodiated adducts.

Peak areas for acyl-chain specific lipid species and standards were quantified by integrating mass spectrometry curves as previously described.[15] For quantitative analyses in cell lines, the peak area was normalized to total protein levels, as well as to the peak area of the corresponding synthetic lipid standard to account for differential loss of samples during lipid extractions and methylation. For analysis of lipid species without a corresponding synthetic lipid standard, the values were normalized to total protein and expressed as normalized peak areas.

### 4. Measurement of Phosphoinositides Using Radioactive $^3$H-inositol Labelling

#### *4a. Culture Conditions and $^3$H-inositol Labelling*

Cells were seeded onto culture dishes pre-coated with 50 μg/mL collagen IV (Thermo Fisher Scientific, cat# 17104019) and cultured in Endothelial Basal Medium (EBM-2) (Lonza, cat# CC- 3156) supplemented with Lonza SingleQuots Kits (cat# CC-4176) and 5% FBS (Thermo Fisher Scientific, cat# 12483020). Just before the cells reached confluence, the growth medium was changed to an inositol-free endothelial cell medium (ScienCell, cat#1001 s.o) supplemented with 75 mCi/mL myo-[2-

$^{3}$H(N)] inositol (Perkin Elmer, cat# NET114A005MC), 5% dialyzed inositol- free FBS, and endothelial growth supplements (ScienCell, cat# 1052) for 24 hours.

***4b. Deacylation and Lipid Extraction***

Following a wash with 1X PBS, adherent cells were scraped and resuspended into 500 µL of 9% perchloric acid and 200 µL of acid washed glass beads in a microcentrifuge tube. The cell suspension was incubated on ice for 5 minutes and then vortexed at top speed for 10 minutes. The sample was then centrifuged at 12,000 × g for 10 minutes at 4°C. The supernatant was carefully discarded and the pellet was resuspended in 1 mL of ice-cold 100 mM EDTA in a water bath sonicator. Once again, the sample was centrifuged at 12,000 × g for 10 minutes at 4°C, and the resulting supernatant was discarded. The pellet was then resuspended in 50 µL of water using a water bath sonicator.

Next, deacylation reagent containing 230 µL of methanol, 130 µL of 40% methylamine, 55 µL of 1-butanol, and 80 µL of water was added to the mixture. The sample was inverted to mix the sample well followed by incubation at room temperature for 20 minutes. The solution was then heated at 53°C for 50 minutes on a heat block, and then dried completely for over 3 hours by vacuum centrifugation. The dried pellet was then resuspended in 300 µL of water by bath sonication and incubated at room temperature for 20 minutes. Again, the sample was dried using vacuum centrifugation for approximately 3 hours. The sample was then resuspended in 450 µL of water using a water bath sonicator and was used as the aqueous phase during extraction.

Lipid extraction reagent containing 240 µL of 1-butanol, 48 µL of ethyl ether, and 12 µL of ethyl formate was added to the aqueous phase. The mixture was vortexed at top speed for 5 minutes and centrifuged at 18,000 × g for 2 minutes to induce phase separation. The lower phase was then carefully transferred to a new microcentrifuge tube. The same extraction procedure was repeated twice by adding the same volumes of lipid extraction reagents to the residual solution. Each time, the lower phase was collected and combined with the others in a microcentrifuge tube. The combined lower phase samples were dried using vacuum centrifugation.

The dried pellet was resolubilized in 50 µL of water followed by water bath sonication. Then, 2 µL of each sample was added to 4 mL of scintillation fluid in a 6 mL polyethylene scintillation vial. Total counts per minute (CPM) emitted by $^{3}$H-inositol incorporated into cellular phosphatidylinositides were quantified with a liquid scintillation using a tritium window. The samples were then stored at -20°C until ready for processing by high performance liquid chromatography (HPLC).

#### *4c. High Performance Liquid Chromatography (HPLC)*

Buffer A was prepared by filtering ultrapure water with a bottle top 0.22 µm vacuum filter, followed by degassing. Buffer B contained 1M ammonium phosphate dibasic ($(NH_4)_2HPO_4$) with pH adjusted to 3.8. Approximately 10 million CPM were loaded into a 2 mL injection vial for a total volume of 55 µL. A blank vial of 55 µL of water was also prepared as a negative control. For HPLC, we used a strong anion exchange (SAX) liquid chromatography column with dimensions 250 x 4.6 mm containing 5 µm silica resin (Phenonenex, Torrance, USA) and equipped with a $^3$H-compatible 500 µL flow cell. Fractionation was done with an Agilent 1200 Infinity series HPLC system controlled by ChemStation software (Agilent, Santa Clara, USA). Flow scintillation of the HPLC eluent was controlled by a BETA-RAM flow detector designed for radio-HPLC, controlled by Laura software (LabLogic, Sheffield, UK).

Following equilibration with Buffers A and B, each sample was automatically injected into the HPLC machine and eluted using an elution profile at a pressure limit of 400 bars, flow rate of 1.0 mL/min and a 90-minute run time. The detection protocol was set up to run for 60 minutes, with a scintillation fluid flow rate of 2.5 mL/min and an 8.57 second dwell time. Briefly, each sample was eluted with 1% Buffer B for 5 min, 1-20% Buffer B for 40 min, 20-100% Buffer B for 10 min, 100% Buffer B for 5 min, 100-1% Buffer B for 20 min, and finally 1% Buffer B for 10 min. At the completion of runs, the HPLC column and flow scintillator were flushed with 100% Buffer A for 30 min at 1.0 mL/min flow rate.

#### *4d. Radioactive Normalization and Analysis*

Using Laura software, all phosphoinositide species were identified based on known retention times. For example, parental phosphatidylinositol (PI) eluted at approximately 10 minutes, PI(3)P at 18 minutes, PI(4)P at 20 minutes, PI(3,5)$P_2$ and PI(3,4)$P_2$ at 29 minutes, and PI(4,5)$P_2$ at 32 minutes. For each species, we recorded the peak area (total counts) and the peak start and end times. For background subtraction, an adjacent region of equal time was subtracted from the total counts of the species. All measured peak areas were normalized against parental PI and against a control condition. Results are expressed as n-fold change compared to control. Data can also be normalized to the total counts if there are significant between group differences in the parental PI levels. However, since parental PI levels are much higher than other species, normalization to total counts did not have a major impact on the data.

### 5. Measuring $PI(4,5)P_2$ Using Fluorescently Labelled Lipid Binding Probes

#### *5a. Culture Conditions and Plasmid Transfection*

Plasmids encoding for the pleckstrin homology (PH) domain of phospholipase delta (PLCδ) fused to GFP to measure $PI(4,5)P_2$ levels were generously gifted from the Grinstein Lab (Hospital for Sick Children, Toronto, Canada).[17] Plasmid transfections were performed as described in Section 2b.

#### *5b. Confocal Microscopy*

Images were acquired using Volocity Software (Perkin Elmer, Woodbridge, Canada) equipped with an Olympus 1X81 confocal microscope, Hamamatsu C9100-13 EM-CCD camera, Yokogawa CSU X1 scanhead, and Improvision Piezo focus drive. All images were analyzed using Volocity software. Briefly, the freehand selection tool was used to draw a region of interest around a segment of the plasma membrane (PM), a cytosolic (Cyt) region excluding the nucleus, and region surrounding the cells as background. Using Volocity software, the mean GFP fluorescence intensity per pixel was measured in the Cyt and PM regions, respectively. The semi-quantitative measure of $PI(4,5)P_2$ at the PM was roughly quantified as a ratio of PM/Cyt GFP fluorescence. Regional background GFP signal measured outside of the targeted cell was subtracted from the PM and Cyt values before computing the PM/Cyt ratio.

### 6. Statistical Analysis

All data represent three independent experiments unless otherwise indicated. Data is represented as mean ± standard deviation, unless otherwise stated. Statistical tests were done using Graphpad Prism 7 Software. * denotes $p<0.05$, ** denotes $p<0.01$, *** denotes $p<0.001$, **** denotes $p<0.0001$, ***** denotes $p<0.00001$.

3. Fontijn, R., Hop, C., Brinkman, H. J., Slater, R., Westerveld, A., van Mourik, J. A. et al.: Maintenance of vascular endothelial cell-specific properties after immortalization with an amphotrophic replication-deficient retrovirus containing human papilloma virus 16 E6/E7 DNA. *Exp Cell Res*

216:199-207, 1995
